# *Bacillus velezensis* 9912 enhances rice growth by triggering plant hormone modulation and complex miRNA-mRNA regulatory networks

**DOI:** 10.1101/2025.03.06.641824

**Authors:** Piao Lei, Yinzhou Jiang, Yan Bai, Jiangchun Hu, Huaqi Pan

**Affiliations:** CAS Key Laboratory of Forest Ecology and Silviculture, Institute of Applied Ecology, Chinese Academy of Sciences, Shenyang 110016, China; University of Chinese Academy of Sciences, Beijing 100049, China

**Author notes:** **Corresponding Authors** (H. Pan).

**Keywords:** plant growth-promoting bacteria, transcriptome, miRNAome, miRNA-mRNA regulatory networks, rice

## Abstract

*Bacillus velezensis*, a prominent member of plant growth-promoting bacteria, effectively suppresses pathogens and promotes plant growth. Despite its well-recognized efficacy, the intricate molecular mechanisms of its growth-promoting attributes remain largely unexplored. In this study, a commercial biopesticide *B. velezensis* 9912 significantly enhanced leaf photosynthesis and improved rice root development in the field experiment. And then, a rigorous hydroponic experiment was performed to investigate its underlying mechanism in plant growth promotion. High-throughput sequencing and bioinformatics analysis revealed 2,938 rice genes and 30 miRNAs responsive to strain 9912. Notably, differentially expressed genes encompassed key transcription factors like ERF, WRKY, and AP2. Remarkably, auxin-and ethylene-related genes were up-regulation significantly at day 1 and pectinesterase-related genes were up-regulation considerably at day 3, which revealed an unusual growth mechanism for strain 9912 to promote rice root development. In addition, DNA replication, cell cycle, cell division, and cytoskeleton organization were significantly enriched in GO and KEGG pathways. Biosynthetic pathways of secondary metabolites, such as carotenoid, lipid, diterpenoid, and brassinosteroid were also significantly enriched. An integrated analysis of the transcriptome and miRNAome identified several miRNA-mRNA regulatory networks, including miR166-Os08g14940, miR396-Os07g9320, miR529-Os12g31540, and miR6249-Os05g07880, which are involved in the process of growth promotion. In conclusion, our investigation offers insightful thoughts into the dynamic responses of rice genes and miRNAs to *B. velezensis* 9912, unveiling plant hormone modulation and potential miRNA-mRNA regulatory networks that are imperative in the growth promotion mechanism.

## 1. Introduction

The global human population is projected to reach approximately 9.7 billion by 2050 (http://esa.un.org/unpd/wpp/). To meet the increasing demand for food, there must be a corresponding expansion in agricultural land or a significant improvement in crop productivity (Tilman et al, 2011). Rice (*Oryza sativa* L.) is among the most significant food crops globally, providing sustenance for nearly 50% of the world′s population (Muthayya et al, 2014). Its extensive cultivation and consumption highlight its significance in addressing global food security challenges. Presently, enhancing rice yields and managing diseases primarily rely on the application of fertilizers and pesticides. However, the excessive use of synthetic chemicals has led to soil degradation, environmental pollution, and potential health risks (Sharma and Singhvi, 2017, Pahalvi et al, 2021).

Plant growth-promoting bacteria (PGPB) are beneficial microorganisms that play a critical role in improving nutrient availability, enhancing plant development, and increasing tolerance to environmental and biological stress (Lugtenberg and Kamilova, 2009, Olanrewaju et al, 2017, Gamalero and Glick, 2011). As a result, the use of potent PGPB is viewed as a promising alternative to synthetic agrochemicals in sustainable agriculture(Vessey, 2003, Atieno et al, 2020). At present, notable reported PGPBs comprise *Bacillus*, *Streptomyces*, *Acinetobacter*, *Bradyrhizobium*, *Klebsiella*, *Pseudomonas*, *Arthrobacter*, *Rhizobium*, *Burkholderia*, and *Aeromonas* (Parray et al, 2016, Vejan et al, 2016, Basu et al, 2021). Among them, *Bacillus* species stand out as a predominant genus, exhibiting traits that promote plant growth through biological processes, such as phytohormone signal transduction, nitrogen utilization, phosphate solubilization, as well as mitigating abiotic and biotic stress (Saxena et al, 2020, Saeid et al, 2018). Hence, research and development of microbial inoculants like *Bacillus* species are necessary and urgent to enhance rice yield and stress resistance.

*Bacillus velezensis* a recently reclassified *Bacillus* species, includes strains such as initial *B. velezensis* strains, *B. methylotrophicus*, *B. amyloliquefaciens* subsp. *plantarum*, and *B. oryzicola* (Ruiz-Garcia et al, 2005, Dunlap et al, 2016, Fan et al, 2017, Adeniji et al, 2019). Research has demonstrated that *B. velezensis* is an exceptional PGPB, exhibiting capabilities in both pathogen resistance and plant growth promotion. For example, *B*. *velezensis* WRN031 significantly enhanced root elongation in maize and rice (Wang et al, 2020), while *B*. *velezensis* NKG-2 improved tomato growth by producing indole-3-acetic acid and siderophores (Harun-Or-Rashid et al, 2018). Additionally, *B*. *velezensis* HNH9 was observed to modify the expression of several growth-related genes, thereby promoting the growth of upland cotton plants (Hasan et al, 2022). In addition to promoting growth, Zhou et al. discovered that *B*. *velezensis* BR-01 demonstrated significant antagonistic effects against various pathogenic fungi and bacteria, including *Magnaporthe oryzae*, *Fusarium fujikuroi*, *Ustilaginoidea virens*, and *Xanthomonas oryzae* pv. *oryza*e (Zhou et al, 2022). These studies demonstrate that *B*. *velezensis* plays a crucial role in enhancing plant growth and pathogen resistance through genes or secondary metabolites associated with growth and defense mechanisms (Fazle Rabbee and Baek, 2020).

Previously, our lab isolated *Bacillus velezensis* strain 9912 from a sedimentary specimen from the Bohai Sea. This strain has become the first commercial biological fungicide in China based on *B. velezensis*, due to its excellent ability to suppress pathogens (Pan et al, 2017). In practical application, it also exhibited outstanding ability to boost the growth of various crops. However, as a typical representative of *B. velezensis*, the growth-promoting mechanism of crop response to strain 9912 remains unclear. In recent years, integrated small RNA sequencing and transcriptome analyses have been applied to detect the responses to light, fertility transition, nitrogen starvation, and zinc deficiency of plants (Lyu et al, 2020, Shin et al, 2018, Zeng et al, 2019, Sun et al, 2021). There is an urgent need to detect the changes in the transcriptome and small RNA of rice after being treated with PGPBs as *Bacillus* spp. In this study, we systematically evaluated the effect of the bioinoculant *B. velezensis* 9912 on the promotion of rice seedling growth in the field experiment. Then we treated rice seedlings with its active ingredient *B. velezensis* 9912 in a hydroponic trial and collected the whole plants to construct mRNA and small RNA libraries for sequencing and bioinformatics analysis. By integrating miRNA and transcriptome data, we explored the molecular mechanisms underlying rice responses to *B. velezensis* treatment, identifying potential miRNA-mRNA interactions involved in the plant’s response.

## 2. Materials and methods

### 2.1. Field experiment design and investigation

The commercial fungicide Aino^®^ Biaobo^®^ with the only active ingredient of *Bacillus velezensis* 9912 came from the North China Pharmaceutical Group Aino Co., Ltd. The field experiment was conducted in the rice field of Ciyutuo Village, Shenbei New District, Shenyang City, Liaoning Province, China (42°04′N, 123°28′E). The microplot area of treated with *B*. *velezensis* 9912 (treatment group) and without *B*. *velezensis* 9912 (control group, CK) were 0.39 and 0.41 acres, respectively. In post-transplantation of rice seedlings, the treatment group was irrigated with Aino^®^ Biaobo^®^ at a rate of 15 kg ha^−1^. After two weeks, samples were collected from the above two microplots using the five-point sampling method in the tillering stage, respectively. Subsequently, plant fresh weight, plant height, stem diameter, average root weight, root length, and lateral root number of the collected samples were measured.

### 2.2. Assessment of chlorophyll and malonaldehyde content in rice seedlings

The total chlorophyll content was performed using a chlorophyll content test kit (BC0990, Solarbio). Approximately 0.1 g of leaves were used for chlorophyll extraction per sample under dark or low-light conditions. According to the test kit instructions, the extract solution (anhydrous ethanol : acetone = 1 : 2, *v*/*v*) was added to the grated tissue and reacted for about 3 h. Absorbance measurements at 645 and 663 nm were conducted by a spectrophotometer. Total chlorophyll content per mg fresh weight was calculated with the formula of (20.21 × A_645_ + 8.02 × A_663_)/fresh weight/1000.

The malonaldehyde (MDA) content was performed using an MDA content assay kit (BC0025, Solarbio). Following the manufacturer’s protocol, approximately 0.1 g of samples mixed with 1 mL of extraction solution were homogenized in an ice bath. Then, the homogenate broth was centrifuged (8000 rpm for 10 min) to obtain supernatant, which was mixed with the working solution and incubated at 100°C for 60 min. Absorbance measurements at 532 and 600 nm were conducted by a spectrophotometer.

### 2.3. Preparation of *B. velezensis* 9912 cells

*B. velezensis* 9912 was cultured on Lysogeny broth (LB) solid medium at 35°C for 48 h. A single colony of the strain was inoculated into 100 mL of LB liquid medium and incubated at 35°C with shaking at 180 rpm for 48 h. The cultured bacterial cells were then transferred into 50 mL sterilized centrifuge tubes and centrifuged for 10 min to pellet the cells. The pelleted cells were resuspended in double-distilled sterile water (ddH2O), and the optical density (OD) was adjusted to 0.5 at 600 nm.

### 2.4. Hydroponic trial to collect RNA nucleic acids

Rice seeds (*Oryza sativa* L. ssp. *japonica* cv. *Nipponbare*) were surface-sterilized by soaking them in 75% (*v*/*v*) ethanol for 10 min. Following sterilization, the seeds were thoroughly rinsed with sterilized water to eliminate any residual ethanol. The sterilized seeds were then placed in a six-well plate for germination and allowed to grow for seven days. Rice seedlings were cultivated under controlled conditions in a plant cultivation rack with an 8-hour dark and 16-hour light cycle at room temperature. For the inoculation of rice seedlings, 5 mL of the *B. velezensis* 9912 cell suspension was applied to the roots of each seedling. As a control, an equal volume of ddH_2_O was applied to the roots of other seedlings. The rice seedlings were watered with ddH_2_O every two days.

### 2.5. mRNA and small RNA libraries preparation

Rice seedlings treated with ddH_2_O and *B. velezensis* 9912, and then collected at day 1 and day 3 were named C1, T1, C3, and T3, respectively. Collected rice samples were stored at - 80℃ refrigerator and delivered to Personalbio company (Personalbio, shanghai, China) for high-throughput sequencing. For the construction of mRNA libraries, the total RNA of rice seedlings was isolated by using the Trizol Reagent. The purity, concentration, and integrity of RNA were assessed with an Agilent Bioanalyzer 2100 (Agilent, Waldbronn, Germany). mRNA enrichment with a poly(A) structure from total RNA was accomplished using Oligo(dT) magnetic beads, and RNA was fragmented into approximately 300 bp fragments via ion disruption. First-strand cDNA synthesis was performed using a 6-base random primer and reverse transcriptase, followed by second-strand cDNA synthesis. To construct small RNA libraries, total RNA underwent ligation to a 3′ adapter and a 5′ adapter using Ligation Enzyme Mix. Subsequently, the samples underwent reverse transcription using Superscript II reverse transcriptase, followed by the amplification of the PCR products. The small RNA libraries were subjected to quality control (QC), with the average insert size measuring approximately 140 to 150 bp. The NEB Next Multiplex Small RNA Library Prep Kit for Illumina was employed for the construction of small RNA libraries. Both the constructed mRNA and small RNA libraries were used for sequencing on the NovaSeq 6000 platform (Illumina) at Personalbio (Personalbio, Shanghai, China.)

### 2.6. Sequencing data quality control, genome mapping, and differential expression analysis

Cutadapt version 1.18 was employed to remove adapters and low-quality reads. The resulting high-quality reads were subsequently mapped to the rice reference genome (Osativa_323_v7.0.fa). HTSeq version 0.9.1 was utilized to calculate the read counts for each gene, which represents the original gene expression levels. Subsequently, FPKM was applied to normalize the expression levels. Regarding raw reads from small RNA libraries, quality information was calculated, and a script developed by Personalbio company was employed to filter the raw data. The resulting clean data were obtained by eliminating adapters and low-quality sequences. Following this, clean reads within a length range of 18 to 36 nt were aligned to the rice reference genome (Osativa_323_v7.0.fa) by miRDeep2 (v2.0.0.8) software. Initially, unique reads were annotated using known miRNAs from the miRBase database. Counting sequences aligned to mature miRNA established their read values. Each miRNA’s read count value was determined by tallying the sequences aligned to the mature miRNA. The chosen miRNA’s abundance for subsequent analysis was selected as the first among those sharing the same name. Finally, the analysis of differential expression for rice genes and miRNAs was conducted utilizing DESeq (v1.38.3). A |log_2_FoldChange| > 1 was established as indicative of a significant expression difference, with a *p*-value threshold of less than 0.05 indicating statistical significance.

### 2.7. Target prediction of differentially expressed (DE) miRNAs

The bidirectional cluster analysis of all miRNAs and samples was carried out by the Pheatmap package (v1.0.12) in the R language. The Euclidean method was employed to calculate distances for hierarchical clustering. Target gene prediction for DE miRNA sequences was conducted using MiRanda (v3.3a) software.

### 2.8. Functional annotation, GO (Gene Ontology) and KEGG (Kyoto Encyclopedia of Genes and Genomes) enrichment analysis

Functional annotation of genes was conducted by BLAST to Swissport and NR databases. Enrichment analyses for differential expression genes (DEGs) and target genes of DE miRNAs were conducted by using GO (http://geneontology.org/) and KEGG (http://www.kegg.jp/). GO enrichment analysis was performed with top GO (v2.50.0), with *p*-values calculated by applying the hypergeometric distribution method. A *p*-value threshold of less than 0.05 was established as the criterion for determining significant enrichment. Significantly enriched GO terms with DEGs were identified to discern the primary biological functions of these genes. Additionally, the clusterProfiler package version 4.6.0 was employed to conduct KEGG pathway enrichment analysis for the target genes of DE microRNAs, focusing on pathways that exhibit significant enrichment (*p*-value < 0.05).

### 2.9. Validation of DGEs by quantitative RT-PCR

Rice genes and miRNAs with distinct expression patterns in each comparison set were randomly selected for qRT-PCR assays. To evaluate the expression of DEGs and DE miRNAs, total RNA was reverse transcribed into cDNA using the PrimerScript First-Strand cDNA Synthesis Kit (Takara, Dalian, China) and miRNA First Strand cDNA Synthesis kit (Sangon Biotech, Shanghai). qRT-PCR experiments were conducted using SYBR Master Premix (Takara, Dalian, China) for DEGs and the miRNA qPCR kit (Sangon Biotech, Shanghai) on a Bio-Rad Real-Time System (BioRad, Hercules, CA, USA). The qRT-PCR conditions for DEGs involved initial steps at 95°C for 2 min, followed by 40 cycles of 95°C for 5 s, 60°C for 30 s, and 65°C for 5 s. For DE miRNAs, the conditions were 95°C for 30 s, followed by 40 cycles at 95°C for 5 s and 60°C for 30 s. Reference genes, including Rice OsACTIN1 (Os03g0718100) and OsUBQ2 (Os02g0161900), along with U6 snRNA, and data were quantified using the 2−ΔΔCt method. Primer synthesis was performed at the Sangon Biotech (Table S1). Each qRT-PCR experiment included three biological replicates and three technical replicates for each sample.

### 2.10. Statistical analysis

Statistically significant differences in plant fresh weight, plant height, stem diameter, average root weight, root length, lateral root number, along with chlorophyll and MDA content of rice treated with and without *B*. *velezensis* 9912 were determined using a *t*-test through SPSS software (SPSS Inc., Chicago, IL, USA). The results were visualized using GraphPad Prism 9 software (GraphPad Inc., San Diego, CA, USA).

## 3. Results

### 3.1. *B. velezensis* 9912 promotes the growth of rice in the field experiment

After two weeks of irrigating the *B*. *velezensis* 9912 (Aino^®^ Biaobo^®^) in the field, the plant fresh weight, plant height, stem diameter, average root weight, root length, lateral root number, along with chlorophyll and MDA content of rice were measured. As shown in Figure 1, the morphological characteristics and physiological responses of rice treated with *B. velezensis* 9912 exhibited a dramatic change compared to the control. The strain 9912 notably enhanced the root length of rice and increased their average root weight and lateral root number. The average root weight of rice treated with *B. velezensis* 9912 exhibited a considerable promotion compared to the control (*p* < 0.05), from 0.31 g to 0.50 g, representing a 60.13% increase. Additionally, the root length was enhanced significantly by 19.30%, from 12.07 cm to 14.40 cm. And lateral root number was also enhanced significantly by 8.51% (Figure 1). Also, a moderate enhancement in the plant fresh weight, plant height, and stem diameter. What should be of concern is the changes in chlorophyll and MDA content. The chlorophyll content was considerably enhanced by 41.06% and the MDA content was decreased by 12.34% (Figure 1).

**Figure 1.**
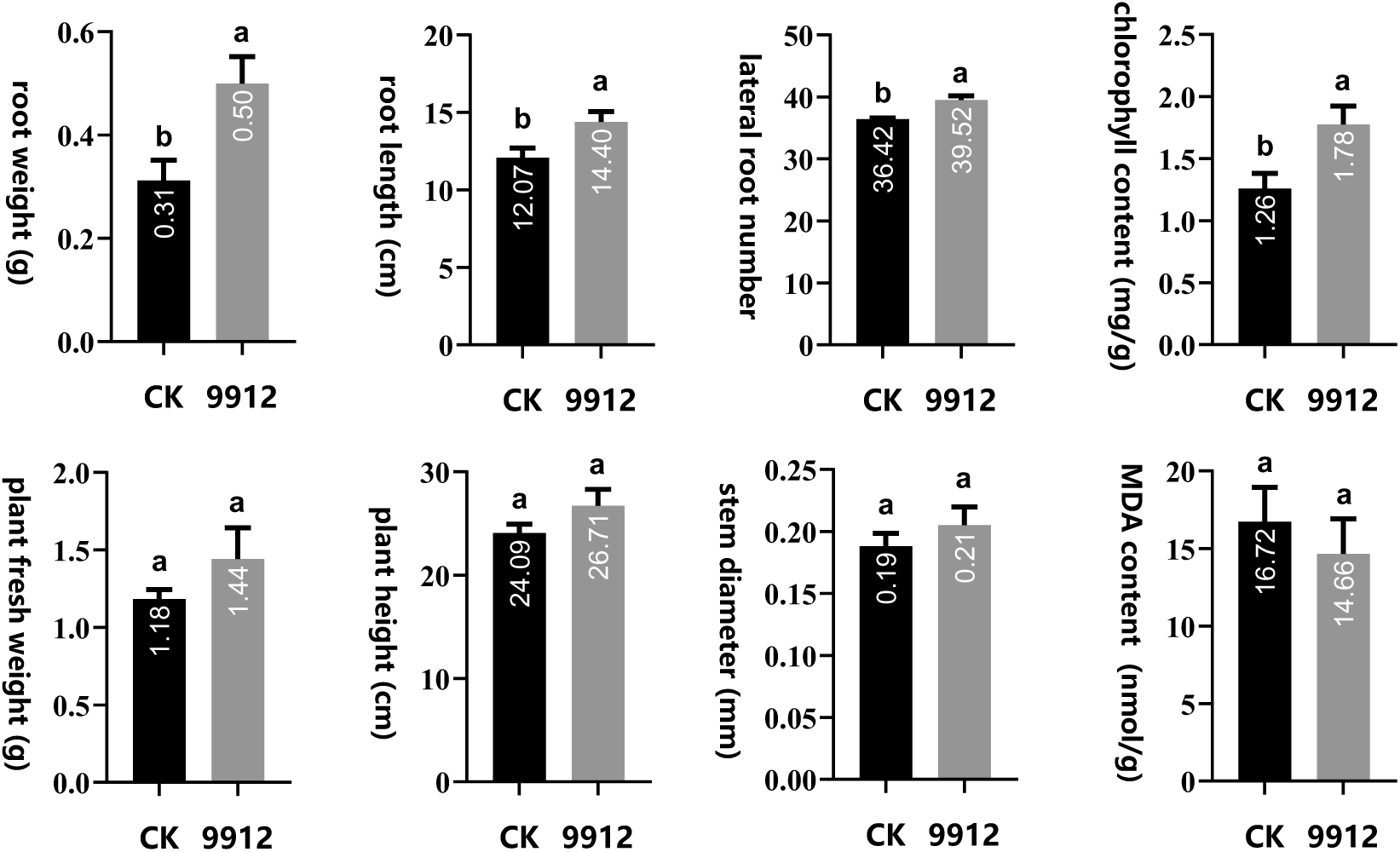
Effects of *B.velezensis* 9912 on promoting the growth of rice seedlings in the field experiment. The *B. velezensis* 9912 (Aino^®^ Biaobo^®^) was used as treatment. The significance of differences between control (CK) and strain 9912 treatment was calculated by *t*-test, significant *p* < 0.05.

### 3.2. Expression of rice genes changed dynamically to *B. velezensis* 9912 treatment

A schematic diagram of transcriptome and small RNA Sequencing of rice seedlings is shown in Figure 2A. Totally, twelve rice samples from hydroponic trials were submitted to Personalbio company (Personalbio, Shanghai, China) for the construction of mRNA libraries for high-throughput sequencing. All twelve mRNA libraries generated 574,547,760 raw reads on the Illumina platform. After quality control, 564,782,960 clean reads were obtained, with Q30 values ranging from 95.33 to 93.27, representing the maximum and minimum values. The subsequent analysis focused on the global transcriptome profiles of rice seedlings using these high-quality clean reads. The majority of clean reads (96.57-98.63%) from each library were successfully aligned to the rice reference genome (Osativa_323_v7.0.fa). HTSeq software and DESeq software were then employed for gene expression calculation and differential expression analysis under the conditions of |log_2_FoldChange| > 1 and *p*-value < 0.05. A total of 2938 rice genes exhibited differential expression across the 12 mRNA libraries.

**Figure 2.**
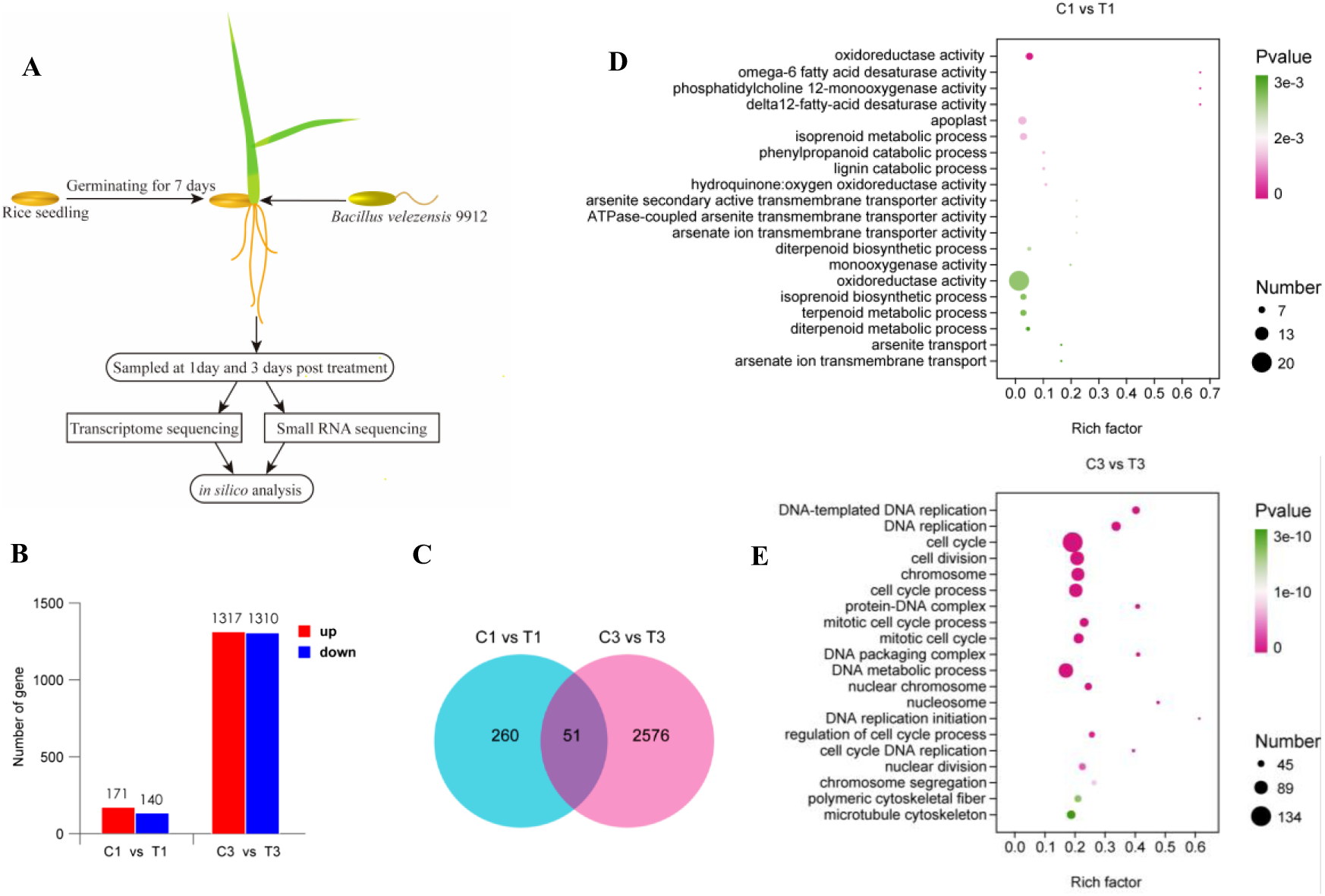
Differentially expressed genes of rice and gene ontology enrichment analysis after *B. velezensis* 9912 treatment. (A) Schematic diagram of transcriptome and small RNA Sequencing of rice seedlings treated by *B. velezensis* 9912. (B) The number of DEGs at day 1 and day 3 post strain 9912 treatment. Rice genes with |log_2_FoldChange| > 1 and significance *p*-value < 0.05 were assigned as considered DEGs. Red bars and blue bars represent the number of up-regulated genes and down-regulated genes, respectively. (C) Venn diagram representing the overlap of rice DEGs after *B. velezensis* 9912 treatment. C1, T1, C3, and T3 represent libraries constructed from rice seedlings treated with ddH_2_O (CK) and *B. velezensis* 9912 and collected at days 1 and 3, respectively. (D) GO enrichment analysis of DEGs in C1 vs T1. (E) GO enrichment analysis of DEGs in C3 vs T3. GO terms were defined as significantly enriched by *p*-value < 0.05. The x-axis represents the rich factor (the number of differential genes annotated to a GO term divided by the total number of genes annotated to that GO term), while the y-axis represents the GO Term. The size of the points in the graph indicates the number of annotated DEGs and the color intensity indicates the significance level.

At day 1 post-treatment with *B. velezensis* 9912, compared with the control, 171 and 140 rice genes were up-regulated and down-regulated, respectively. At day 3 post-treatment, rice genes responded more dramatically to *B. velezensis* 9912. A total of 2627 DEGs were identified, comprising 1317 upregulated genes and 1310 downregulated genes in rice (Figure 2B). Among these DEGs, 51 genes exhibited differential expression in mRNA libraries collected at both day 1 and day 3 (Figure 2C and Table S2). Notably, several genes, such as Os02g51440, Os07g44550, and Os04g55740, were up-regulated in both C1 *vs* T1 and C3 *vs* T3, while Os01g48446, Os09g35020, and Os09g35010 were down-regulated in both comparisons (Table S2). Most DEGs displayed distinct expression trends at two timepoints. For instance, the log_2_(FoldChange) values for Os02g36190 were -1.84 in C1 *vs* T1 and 3.36 in C3 *vs* T3, respectively (Table S2). We performed qRT-PCR to validate the expression patterns of DEGs in response to the treatment with *B. velezensis* 9912. Three genes were randomly selected for validation: Os10g31540, Os08g24300, and Os06g03380. The values of log_2_FoldChange of these genes were 4.10, 2.51, and -2.13, respectively. The relative expression levels of these genes in response to treatment with *B. velezensis* 9912 were 2.48, 1.35, and 0.29 times that of the control group, respectively (Table S2 and Figure S1). The regulation trends of these DEGs were consistent with *in silico* analysis.

### 3.3. DEGs were related to growth regulation

We conducted annotation of DEGs by performing BLAST searches against the Swiss-Prot and NR databases. The results revealed specific DEGs identified in C1 *vs* T1, such as Os10g31540, Os07g43260, Os01g67410, Os08g44960, and Os03g43400, which encode cell wall structural protein, SKP1-like protein, AP2-like ethylene-responsive transcription factor, ERF transcription factor, and auxin-responsive protein, respectively (Table 1). In C3 *vs* T3, DEGs included Os12g31540, Os01g43851, Os03g50420, and Os01g20980, coding for MID1-like protein, cytochrome P450, cell division-associated protein, and pectinesterase/pectinesterase inhibitor, respectively (Table 1). In addition, we observed that 18 and 208 DEGs, which encode leucine-rich repeat receptors, were identified in the C1 *vs* T1 and C3 *vs* T3 comparisons, respectively. Additionally, 9 and 24 DEGs were annotated as encoding peroxidases, while 3 and 2 DEGs were noted to encode laccases in the C1 *vs* T1 and C3 *vs* T3 comparisons, respectively (Table S2). These results suggested that the inoculation with *B. velezensis* 9912 altered the expression of leucine-rich repeat receptors, which may lead to cascading reactions in the form of reactive oxygen species burst and other disease-resistant immune responses.

**Table 1.**
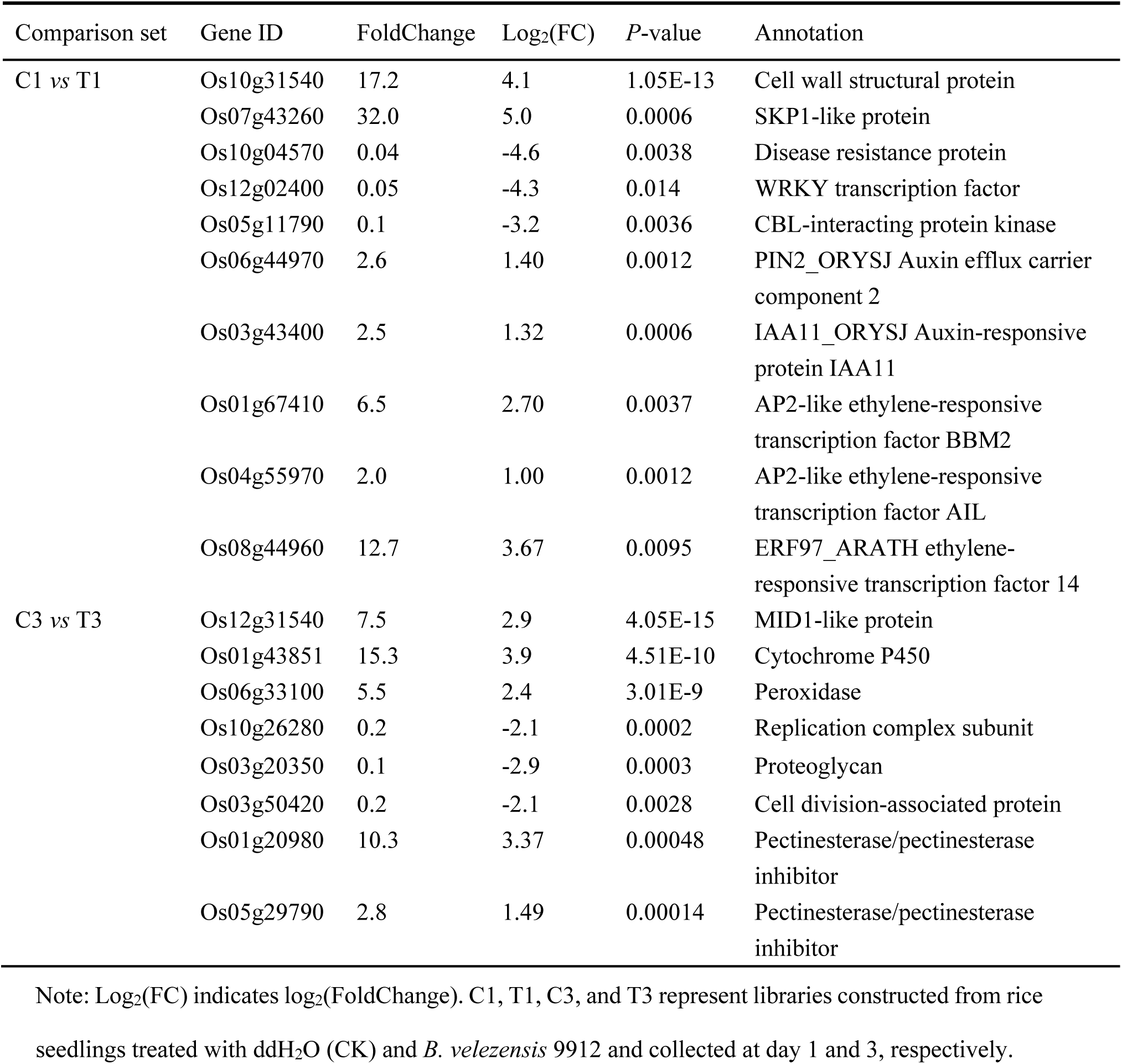
Function annotation of selected differentially expressed genes post *B. velezensis* 9912 treatment.

Furthermore, we conducted GO and KEGG enrichment analyses to explore the pathways involving DEGs. In C1 *vs* T1, significantly enriched GO terms were related to oxidoreductase activity, isoprenoid metabolic processes, and omega-6 fatty acid desaturase activity. Additionally, significantly enriched KEGG pathways were associated with phenylpropanoid biosynthesis, diterpenoid biosynthesis, and carotenoid biosynthesis (Figures 2D and S2A). In C3 *vs* T3, the top significant GO terms were enriched in DNA replication, cell division, and cell cycle. KEGG pathways were associated with diterpenoid biosynthesis, DNA replication, glutathione biosynthesis, and zeatin biosynthesis (Figures 2E and S2B). These findings indicated that rice genes related to growth regulation were responsive to *B. velezensis* 9912 at day 1 and day 3 post-treatment. DEGs participate in pathways linked with DNA replication, cell cycle, and other secondary metabolites, which require transcription factors for initiation. We identified the types of transcription factors to which DEGs belonged. At day 1 post-treatment, DEGs belonged to ERF, WRKY, and AP2 (Figure 3A), while bHLH, WRKY, and MYB were the top three transcription factors at day 3 post-treatment (Figure 3B). It is noteworthy that most transcription factors were down-regulated after *B. velezensis* 9912 inoculation (Figure 3).

**Figure 3.**
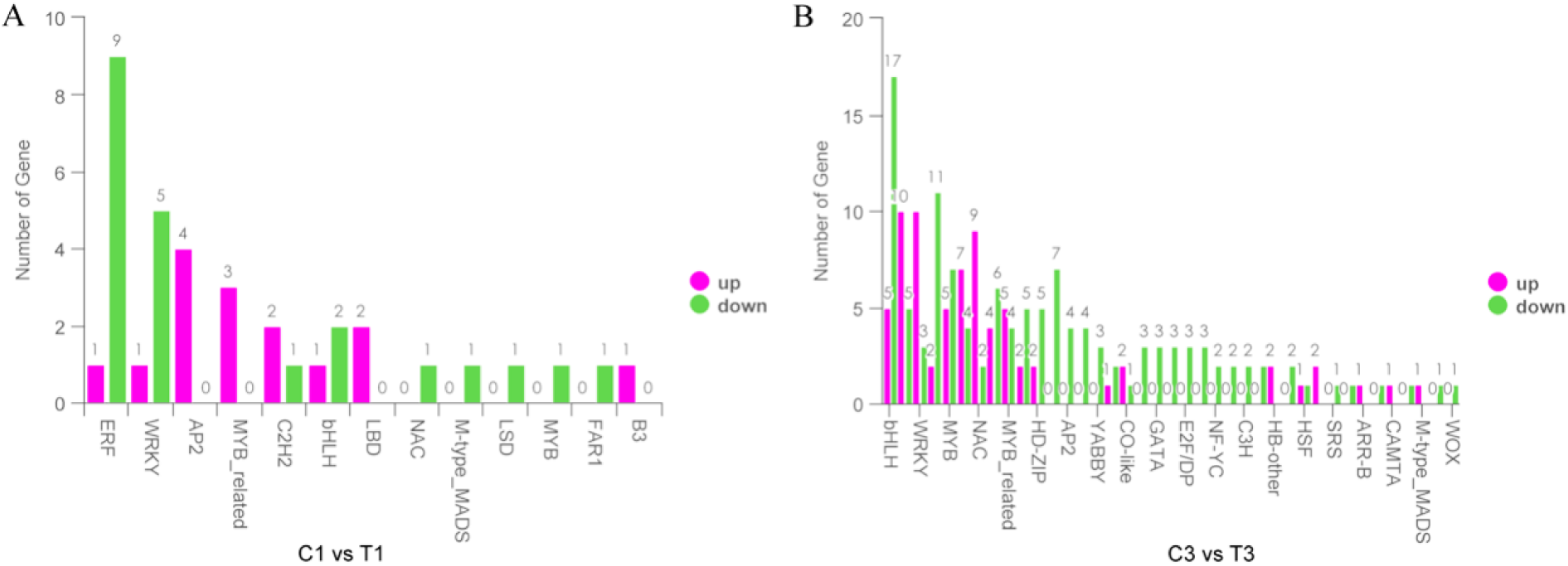
Distribution of differentially expressed transcription factors. (A) Differentially expressed transcription factors of rice seedlings at day 1 post *Bacillus velezensis* 9912 treatment. (B) Differentially expressed transcription factors of rice seedlings at day 3 post *B. velezensis* 9912 treatment. The x-axis represents different transcription factor families, while the y-axis represents the number of differential genes assigned to each transcription factor family. Purple bars and green bars represent the number of up-regulated transcription factors and down-regulated transcription factors, respectively. C1, T1, C3, and T3 represent libraries constructed from rice seedlings treated with ddH_2_O (CK) and *B. velezensis* 9912 and collected at days 1 and 3, respectively.

### 3.4. Rice miRNAs were responsive to *B. velezensis* 9912 treatment

Small RNA sequencing produced a total of 543,152,528 raw reads, with 167,815,966 clean reads remaining after quality control. Utilizing BLAST, unique reads were aligned against the Rfam13 database to identify known non-coding RNAs (ncRNAs), including rRNA, tRNA, snRNA, and snoRNA. Reads aligning with these ncRNA categories were excluded, and the remaining reads proceeded to the next step of the analysis. The number of miRNA sequences after deduplication using unique reads aligned to the Rfam13 database ranged from 2721 to 5249. Furthermore, by blasting against the miRBase databases, 482 known miRNAs were detected in the 12 small RNA libraries, with the number of expressed miRNAs in each library ranging from 302 to 367. The DESeq software was utilized to identify differentially expressed (DE) miRNAs under the condition of |log_2_FoldChange| > 1 and a significance *p*-value < 0.05. In total, thirty miRNAs were detected as differentially expressed in all small RNA libraries. At day 1 post-treatment *B. velezensis* 9912, only two miRNAs, osa-miR5799 and osa-miR821c were up-regulated, with no down-regulated miRNAs identified (Table S3). Meanwhile, at day 3 post-treatment, nine miRNAs were up-regulated and nineteen miRNAs were down-regulated (Table S3). And we performed qRT-PCR to validate the expression trends of DE miRNAs Osa-miR166e and Osa-miR319b were selected for qRT-PCR validation. The relative expression levels of all the selected DE miRNAs observed through qPCR were consistent with the results obtained from high-throughput sequencing (Figure S1 and Table S3). Additionally, hierarchical clustering analysis showed that the expression trends of DE miRNAs were categorized into six primary clades. For example, osa-miR1440b, osa-miR395y, and osa-miR827 shared an expression trend of low expression in C1 and C3 libraries, while a high expression in T1 and T3 libraries (Figure S3). Osa-miR1423-3p, osa-miR171h, and osa-miR186I expressed with a trend of high expression in C1 and T1 libraries, and low expression in C3 and T3 libraries (Figure S3). These results showed the potential trends of DE miRNAs in rice at different growth times, stimulated with or without strain 9912.

### 3.5. Potential target genes of DE miRNAs involved in cellular biosynthesis

To unravel the potential biological functions of miRNAs responsive to *B. velezensis* 9912, psRobot was employed to predict miRNA-targeted genes, revealing a total of 257 potential target genes for DE miRNAs. For instance, osa-miR821c was predicted to target Os02g38340, encoding an actin-like ATPase protein, while osa-miR5799 targeted Os03g24730, Os12g38210, and Os10g32050, which code for galactose oxidase, plant U-box protein, and ankyrin repeat protein, respectively. Furthermore, miR166e was predicted to target Os08g14940, encoding a leucine-rich repeat protein, and osa-miR3981-5p targeted a terpene synthase gene, Os03g22634. Additional genes such as Os11g24484, Os02g56370, and Os09g02360, coding for beta-ketoacyl reductase, wall-associated kinase, and cyclin, respectively, were potentially targeted by osa-miR2867-3p, osa-miR827, and osa-miR529a (Table S4).

Subsequently, we conducted GO and KEGG enrichment analysis of potential target genes of DE miRNAs. One day post *B. velezensis* 9912 treatment, the most significantly enriched GO terms for cell components were Arp2/3 protein complex (GO:0005885), actin cytoskeleton (GO:0015629), and cytoskeleton (GO:0005856) (Figure 4A). In terms of molecular function, the top enriched terms included cysteine-type deubiquitinase activity (GO:0004843), ubiquitin-like peptidase activity (GO:0019783), and deubiquitinase activity (GO:0101005) (Figure 4A). Additionally, the foremost enriched biological processes were linked to actin nucleation (GO:0034314 and GO:0045010) and actin cytoskeleton organization (GO:0007015 and GO:0030029) (Figure 4A). Conversely, at day 3 post *B. velezensis* 9912 treatment, enriched GO terms were associated with the membrane (GO:0016020, GO:0008250, and GO:0005741), copper ion binding (GO:0005507), catalytic activity (GO:0003824), GMP biosynthetic/metabolic processes (GO:0006177/GO:0046037), and endocytosis (GO:0006897) (Figure 4B). Moreover, KEGG enrichment analysis of potential target genes corresponding to DE miRNAs indicated significant enrichment in pathways such as endocytosis, carotenoid biosynthesis, ether lipid metabolism, glycerophospholipid metabolism, brassinosteroid biosynthesis, and plant-pathogen interaction (Figure S4).

**Figure 4.**
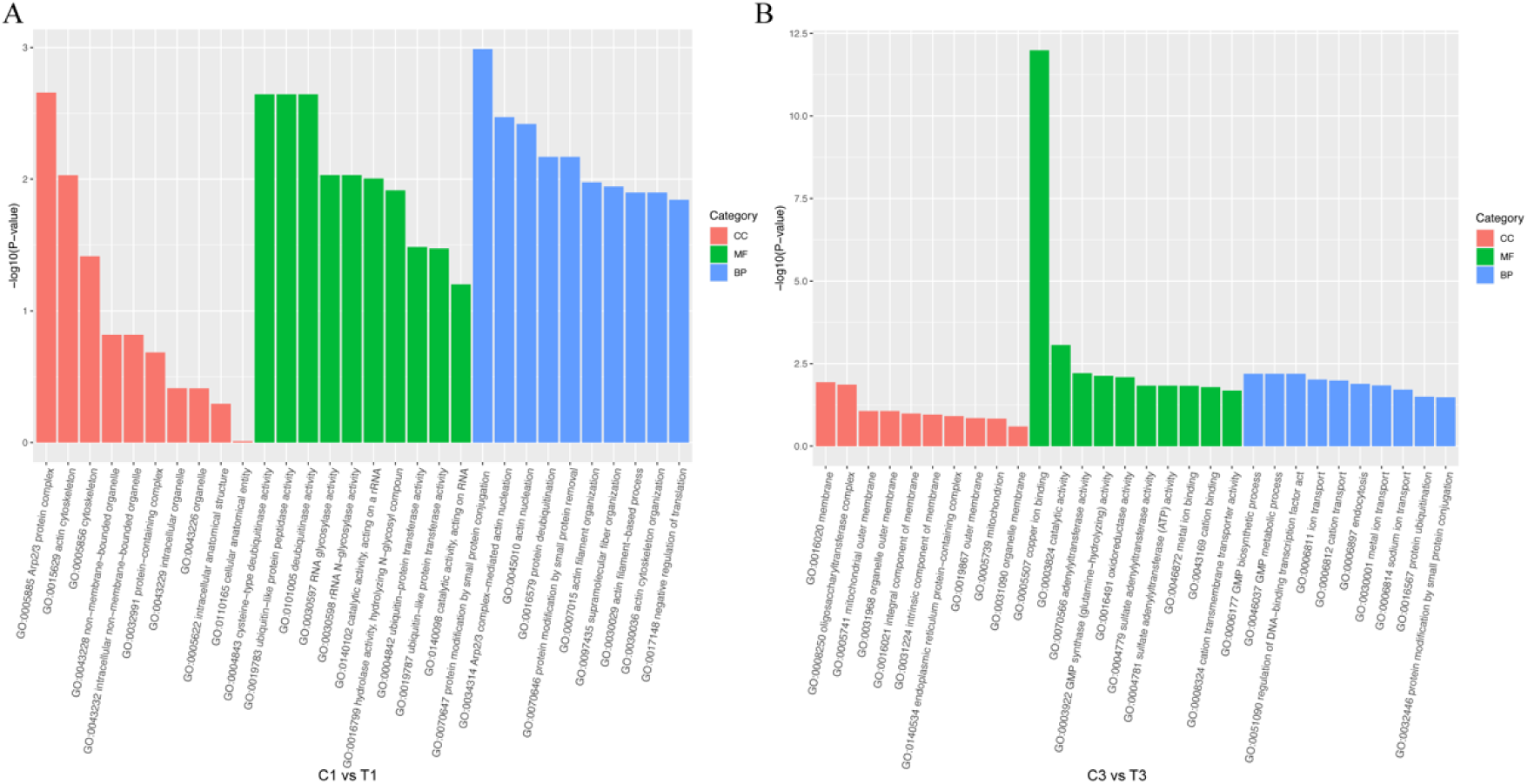
Gene ontology enrichment analysis of target genes of differentially expressed miRNAs post *Bacillus velezensis* 9912 treatment. (A) GO enrichment analysis of target genes of DE miRNAs in C1 vs T1. (B) GO enrichment analysis of target genes of DE miRNAs in C3 vs T3. C1, T1, C3, and T3 represent libraries constructed from rice seedlings treated with ddH2O (CK) and *B. velezensis* 9912 and collected at days 1 and 3, respectively. The X-axis represents the GO level 2 terms, and the Y-axis represents the enrichment -log10 (*p*-value) for each term. GO enrichment analysis results of the target genes of DE miRNAs were categorized into Molecular Function (MF), Biological Process (BP), and Cellular Component (CC). The top ten significantly enriched GO terms were shown in GO enrichment results.

### 3.6. *B. velezensis* 9912 regulated rice growth through miRNA-mRNA networks

Differential expression analyses of both transcriptome and small RNA sequencing unveiled a myriad of DEGs and DE miRNAs in response to *B. velezensis* 9912 treatment compared to the control. Notably, a comprehensive joint analysis was conducted to elucidate regulatory patterns involving DE miRNAs and genes post *B. velezensis* 9912 treatment. As shown in Figure 5, The networks of DE miRNAs and genes were grouped into 29 nodes arranged in 23 clusters with ≥ 3 nodes per cluster, and 6 with two nodes. There were seven main DE miRNAs (≥ 10 nodes) in the network, including Osa-miR6249b (Figure 5A), Osa-miR529a (Figure 5B), Osa-miR396b-5p (Figure 5C), Osa-miR5799 (Figure 5D), Osa-miR166e-5p (Figure 5E), Osa-miR437, and Osa-miR397b.

**Figure 5.**
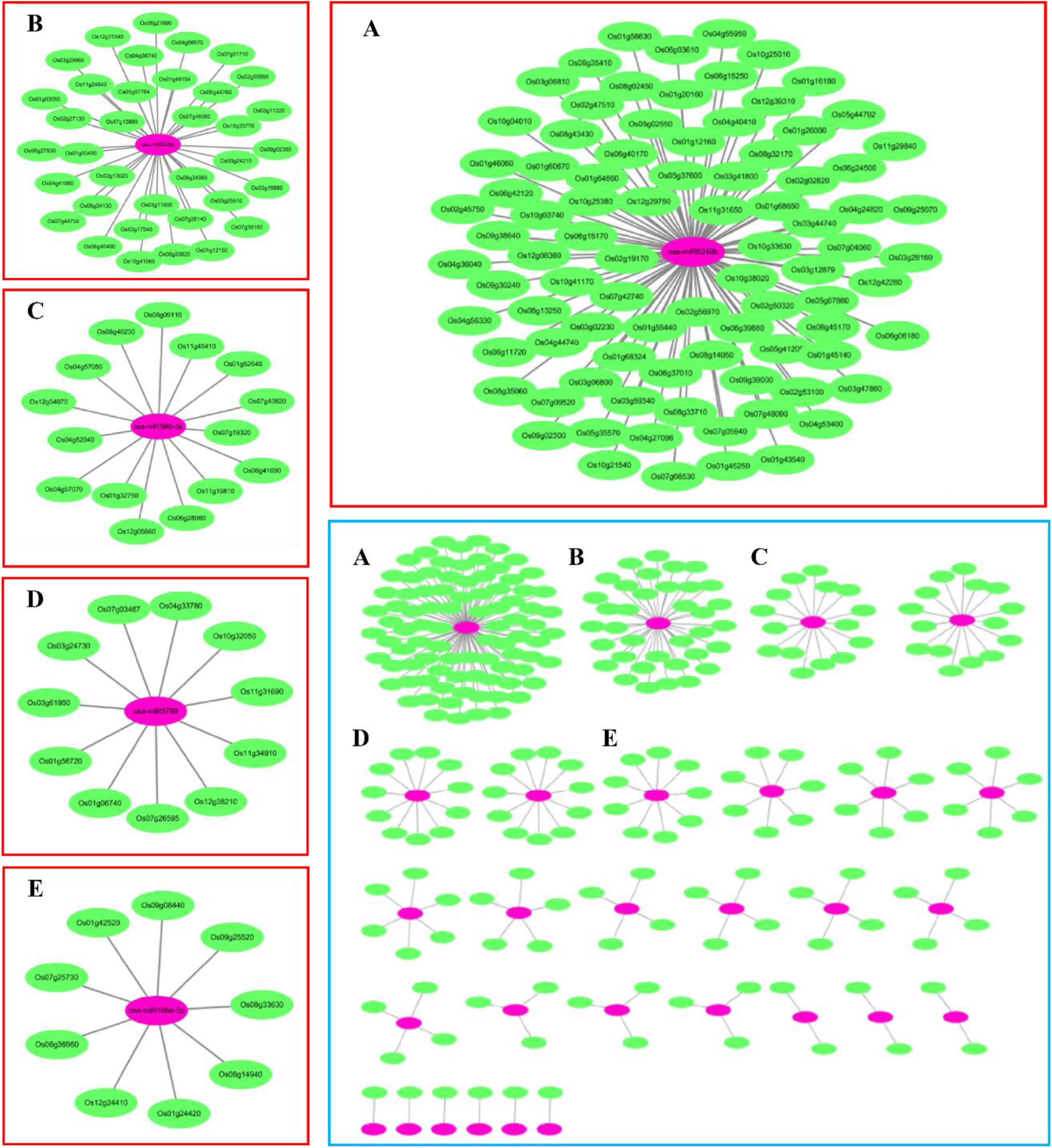
Network maps of the regulatory target genes of differentially expressed miRNAs. Regulatory networks were developed by Cytoscape v3.6.1. miRNAs are shown as purple nodes, Gene IDs are shown as green nodes.

Further analysis indicated that certain DE miRNAs orchestrated the regulation of their target genes, thereby promoting rice growth. For instance, Osa-miR5799 targeted Os01g06740, which encodes RNA glycosylase involved in macromolecule metabolism. Osa-miR166e-5p was predicted to target Os08g14940, a receptor kinase implicated in signal transduction (Figures 5 and 6). Osa-miR396b-5p targeted a PR protein, Os07g9320, influencing rice disease resistance (Figures 5 and 6). Additionally, Osa-miR6249b and Osa-miR529a were predicted to target Os05g07880 and Os12g31540, associated with phospholipase and encoding a MID1-like protein, respectively, both contributing to the regulation of cell number and lipid metabolism (Figures 5 and 6).

**Figure 6.**
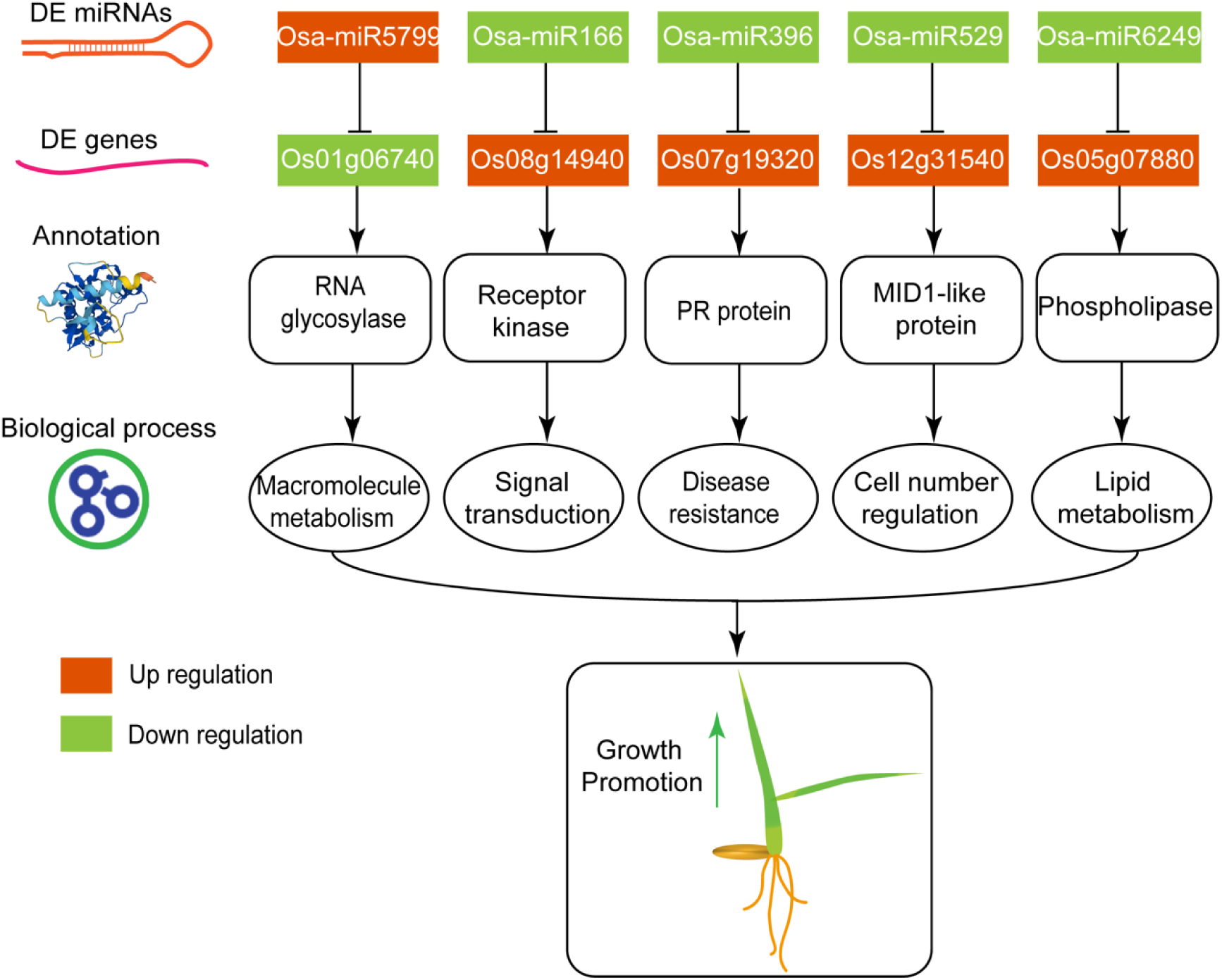
Potential regulatory network between differentially expressed miRNAs and genes that promote the growth of rice seedlings after *B. velezensis* 9912 treatment. Orange color and green color represent upregulation and downregulation, respectively.

## 4. Discussion

*Bacillus* spp. have been extensively investigated as PGPB, renowned for their capacity to growth promotion and ameliorate abiotic and biotic stress tolerance (Saeid et al, 2018, Saxena et al, 2020). Numerous studies have documented the growth-promoting effects of *Bacillus* on rice. Chung et al. observed a substantial increase in shoot length, rising from 11.67 cm to 19.33 cm during the seedling stage in test tubes, following the application of a *B. oryzicola* YC7007 suspension to the rice rhizosphere (Chung et al, 2015). Meanwhile, *B. cereus* YN917 was found to significantly boost the growth of rice. After a 30-day inoculation period, rice seedlings treated with YN917 exhibited substantial improvements in various growth parameters compared to the control, including plant height (from 22.65 cm to 33.69 cm), root length (2.98 cm to 7.68 cm), and fresh weight (from 1.75 g to 2.28 g) (Zhou et al, 2021). In this investigation, rice treated with *B. velezensis* 9912 exhibited notable changes in morphological characteristics compared to the control in the field (Figure 1), including a significant enhancement in the average root weight, root length, and lateral root number. In addition, chlorophyll also increased significantly, suggesting that strain 9912 promoted rice growth by enhancing photosynthesis and promoting root development. The significant decrease in MDA suggested that 9912 could improve the environmental resistance of rice by scavenging oxygen-free radicals.

Although these *Bacillus* spp. can significantly promote root development in rice, the growth mechanism of rice roots remains largely unexplored. Here, the analysis of DEGs related to growth regulation, in C1 *vs* T1, auxin efflux carrier PIN-related gene (Os06g44970), auxin-responsive protein IAA11-related gene (Os03g43400), along with ethylene-responsive transcription factor BBM2 and AIL-related genes (Os01g67410 and Os04g55970) were up-regulation significantly (Table 1), which suggested the inoculation with *B. velezensis* 9912 increased the secretion of auxin or ethylene to promote root growth. As is well-known, pectinesterase (PME), as an important pectin enzyme, is essential in regulating seed germination and root extension (Wang et al, 2022). In C3 *vs* T3, pectinesterase-related genes (Os01g20980 and Os05g29790) were up-regulation significantly (Table 1), which indicated the inoculation with *B. velezensis* 9912 increased the accumulation of PME to promote root growth. These changes of DEGs were consistent with the changes in morphological characteristics. Furthermore, these results suggested *B. velezensis* 9912 had a remarkable capacity to promote the root growth of rice by stimulating the plant hormone modulation at the initial stages, subsequently enhancing the expression of PME. These findings contribute to a more comprehensive understanding that PGPB promotes crop growth, moving beyond the commonly believed that *Bacillus* spp. secrete auxin for crop growth (Saxena et al, 2020, Saeid et al, 2018).

Moreover, we have found that most DE miRNAs are conserved miRNAs. Such as miR166, miR171, miR319, miR396, miR437, miR529, and miR821 differentially expressed at day 1 and day 3 post *B. velezensis* 9912 treatment. It is important to note that conserved miRNAs and their target genes play pivotal roles in plant development, nutrient utilization, and responses to environmental stimuli (Table S4). Besides, *Bacillus* spp. also alter the expression of miRNAs of plants against pathogens. After inoculation of *B. velezensis* FZB42, defense-related miRNAs, including 4 novel miRNAs differentially expressed (miR159, miR400, and miR8167 exhibited the highest changes) and 11 known miRNAs (Xie et al, 2017). In wheat treated with *B*. *subtilis* 26D, the expression of miR156, miR160, miR166a, and miR398 involved in phytohormone signaling pathways changed dramatically, which may help against phloem-feeding insects (Rumyantsev et al, 2023). Here, we describe the first miRNAome of rice response to *Bacillus* PGPB and shed light on the growth-promoting mechanism of *Bacillus* sp. by triggering miRNA-mRNA regulatory networks of rice.

The joint analysis of transcriptome and small RNAome is crucial for a comprehensive understanding of gene regulation in plants, unraveling intricate networks governing developmental processes, stress responses, and environmental adaptations (Li et al, 2022, Li et al, 2017, Lyu et al, 2020). Several studies have utilized a joint analysis of the transcriptome and small RNAome to elucidate the molecular mechanisms through which plants respond to biotic stress, abiotic stimuli, and other biological processes. Li et al. discovered *B. amyloliquefaciens* LZ04 enhances the resistance of *Arabidopsis thaliana* to high calcium stress. Their study revealed the existence of a sophisticated regulatory network involving 23 lncRNAs, 10 miRNAs, and 80 mRNAs, among which miR159c, miR164a, miR172e, and their corresponding target genes played pivotal roles (Li et al, 2020). Sun et al identified 13 distinct pairs of miRNA-mRNA with notable expression differences, including miR156, miR5488, miR399 and their potential target genes, suggesting potential roles of miRNA-mRNA regulatory networks involved in regulating fertility changes in rice (Sun et al, 2021). Through integrated analysis of miRNAome and transcriptome, we further identified miRNAs and their potential target genes that regulated rice seedling′s growth. Such as miR166-Os08g14940 (receptor kinase), miR396-Os07g9320 (PR protein), miR529-Os12g31540 (MID1-like protein), and miR6249-Os05g07880 (phospholipase) (Figure 6). Using miR166 and miR396 as examples, in rice, miR166 plays a crucial role in nutrient uptake, shoot meristem initiation, and leaf cell development (Iwamoto, 2022, Nagasaki et al, 2007, Li et al, 2019). Additionally, miR396 is known to modulate grain size, shoot architecture, as well as balance growth and resistance to pathogens(Miao et al, 2020, Chandran et al, 2019). These miRNA-mRNA regulatory networks indicated the growth-promoting mechanism of the strain 9912 at the molecular level.

In this investigation, GO and KEGG enrichment analysis indicated that DEGs were linked to DNA replication, cell division, and cell cycle (Figures 2 and S2). These results were consistent with GO and KEGG enrichment of target genes of DE miRNAs (Figures S3 and S4). Moreover, our present study demonstrated that second metabolites biosynthesis pathways were also enriched, such as phenylpropanoid biosynthesis, diterpenoid biosynthesis, and carotenoid biosynthesis (Figure 2). Other studies also found DE miRNA-mRNA regulatory networks were related to the lignin synthesis of anther walls, flavonoid metabolism pathway, and other secondary metabolites and biosynthesis/metabolic pathways during development and response to pathogen (Sun et al, 2021, Zhao et al, 2020). In addition, we found that DEGs were related to signal transduction, peroxidase, and pathogenesis-related (PR) protein, which are considered *B. velezensis* 9912 to induce disease resistance in rice seedlings.

In addition, during the harvest period, rice increased yield by equivalent to 5% under inoculation with strain 9912 in the field (data not shown). And previous studies in our lab indicated that *B. velezensis* 9912 exhibited antagonistic activity in controlling a range of plant pathogens (Pan et al, 2017). Combining these above results and our findings in the present study, microbial inoculant *B. velezensis* 9912 improves rice growth and health through multiple pathways, including phytohormone signal transduction, miRNA-mRNA regulatory networks, induced systemic resistance, as well as mitigating pathogen invasion. Thus, the *B. velezensis* 9912 may play an important role in the sustainable production of rice.

In conclusion, *B. velezensis* 9912 significantly promoted the growth of rice in the field experiment by enhancing leaf photosynthesis and improving root development. Transcriptome and miRNAome analysis identified 2938 DE rice genes and 30 miRNAs responsive to *B. velezensis* 9912 treatment. DEGs were associated with DNA replication, cell cycle, cell division, and secondary metabolites biosynthesis/metabolism pathways. Notably, auxin- and ethylene-related genes were up-regulation significantly at day 1 and then pectinesterase-related genes were up-regulation significantly at day 3, which suggested *B. velezensis* 9912 stimulates rice biosynthesis of ethylene and auxin at the initial stages, subsequently enhancing the expression of PME to promote root growth. Furthermore, we described the first miRNAome and miRNA-mRNA networks of rice response to the genus *Bacillus*. And revealed miRNA-mRNA regulatory networks that regulate the growth of rice after *B. velezensis* 9912 treatment, including key players like miR166-Os08g14940, miR396-Os07g9320, miR529-Os12g31540, and miR6249-Os05g07880. The findings presented above establish a theoretical foundation for further investigation into the molecular mechanisms underlying the growth promotion induced by *B*. *velezensis* 9912 and provide valuable insights into the dynamic responses of rice genes and miRNAs to PGPBs.

## Acknowledgments

This study was supported by the National Key Research and Development Program of China (2023YFD1501200 and 2023YFD1500500), the Strategic Priority Research Program of the Chinese Academy of Sciences (XDA28090300), the Liaoning Province Applied Basic Research Program (2022JH2/101300185 and 2022JH2/101300186), and the Youth Innovation Promotion Association CAS (Y2022063).

## Support Information

**Figure S1.**
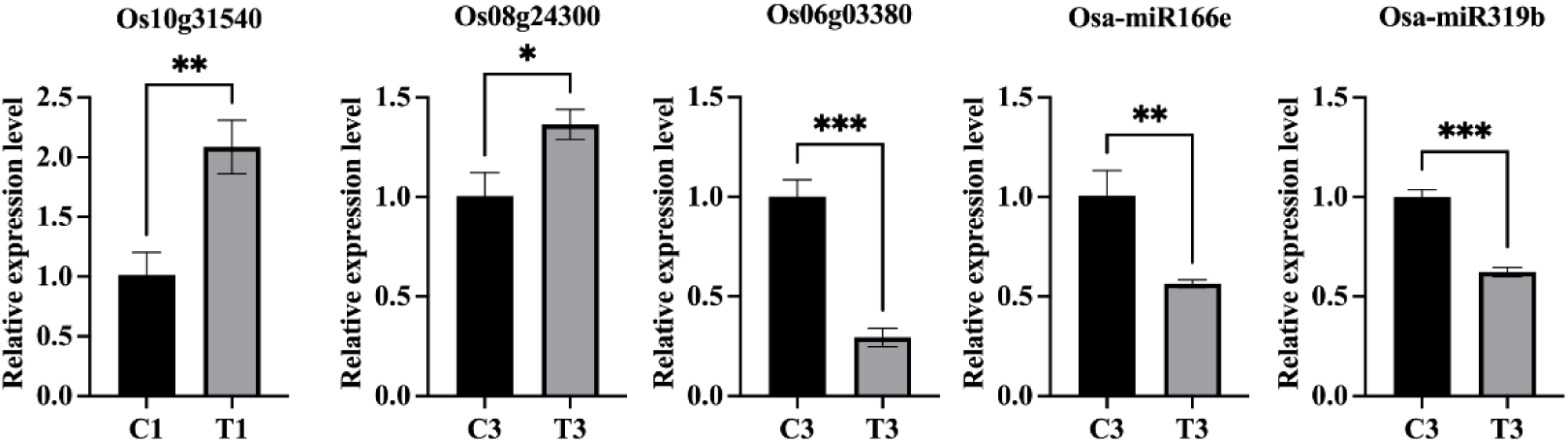
qRT-PCR validates the expression trends of differentially expressed genes and miRNAs identified in high throughput sequencing. C1, T1, C3, and T3 represent libraries constructed from rice seedlings treated with ddH2O (CK) and *B. velezensis* 9912 and collected at 1 and 3 days, respectively. The significance of differences between control (CK) and *B. velezensis* 9912 treatment was calculated by t-test, “ns” and asterisks represent no significant difference and significant difference at *p* < 0.05, respectively.

**Figure S2.**
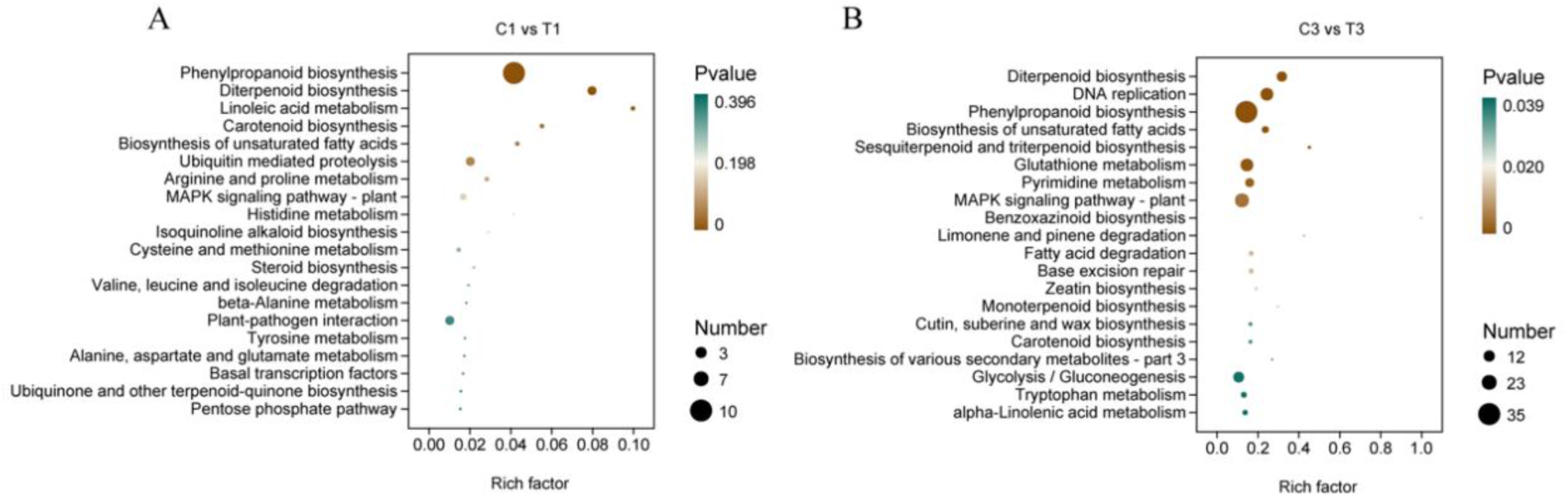
KEGG enrichment analysis of differentially expressed genes post *Bacillus velezensis* 9912 treatment. (A) KEGG enrichment analysis of differentially expressed genes in C1 vs T1. (B) KEGG enrichment analysis of target genes of differentially expressed miRNAs in C3 vs T3. Clusterprofiler was used for was applied to KEGG enrichment analysis. Gene lists and the number of genes for each KEGG pathway were calculated by using differentially expressed genes annotated with KEGG pathway information. Significant enrichment threshold set at *p*-value < 0.05. The X-axis represents the rich factor (the number of differential genes annotated to a pathway divided by the total number of genes annotated to that KEGG pathway), while the Y-axis represents the KEGG pathway. The size of the points in the graph indicates the number of annotated differentially expressed genes and the color intensity indicates the significance level. C1, T1, C3, and T3 represent libraries constructed from rice seedlings treated with ddH_2_O (CK) and *B. velezensis* 9912 and collected at 1 and 3 days, respectively.

**Figure S3.**
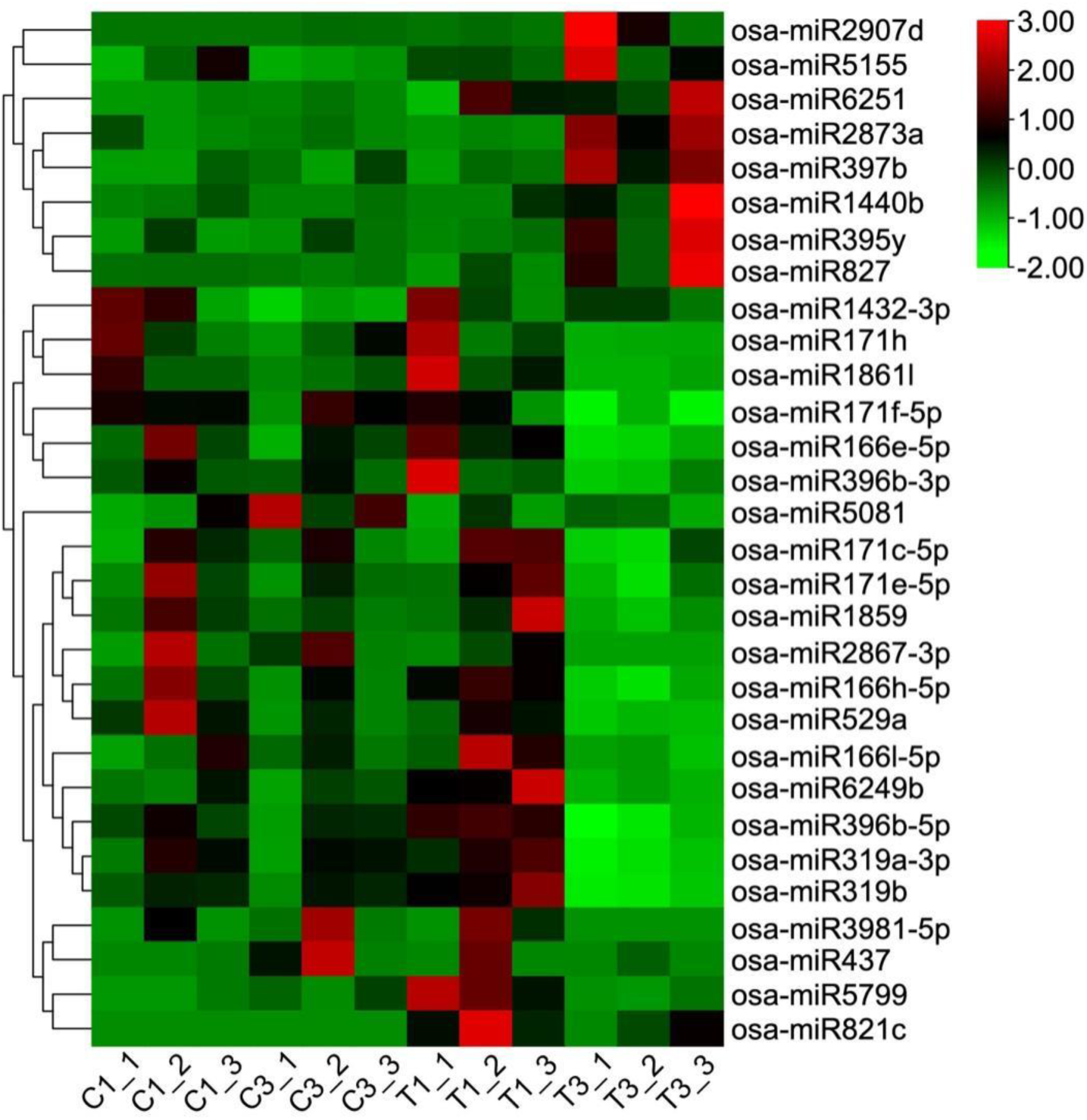
Clustering heatmap of differentially expressed rice miRNAs. Each column represents a sample, and the rows represent miRNA genes. Hierarchical clustering was performed using the Euclidean and complete linkage method. Red indicates high miRNA expression, while green indicates low miRNA expression. C1, T1, C3, and T3 represent libraries constructed from rice seedlings treated with ddH_2_O (CK) and *B. velezensis* 9912 and collected at days 1 and 3, respectively.

**Figure S4.**
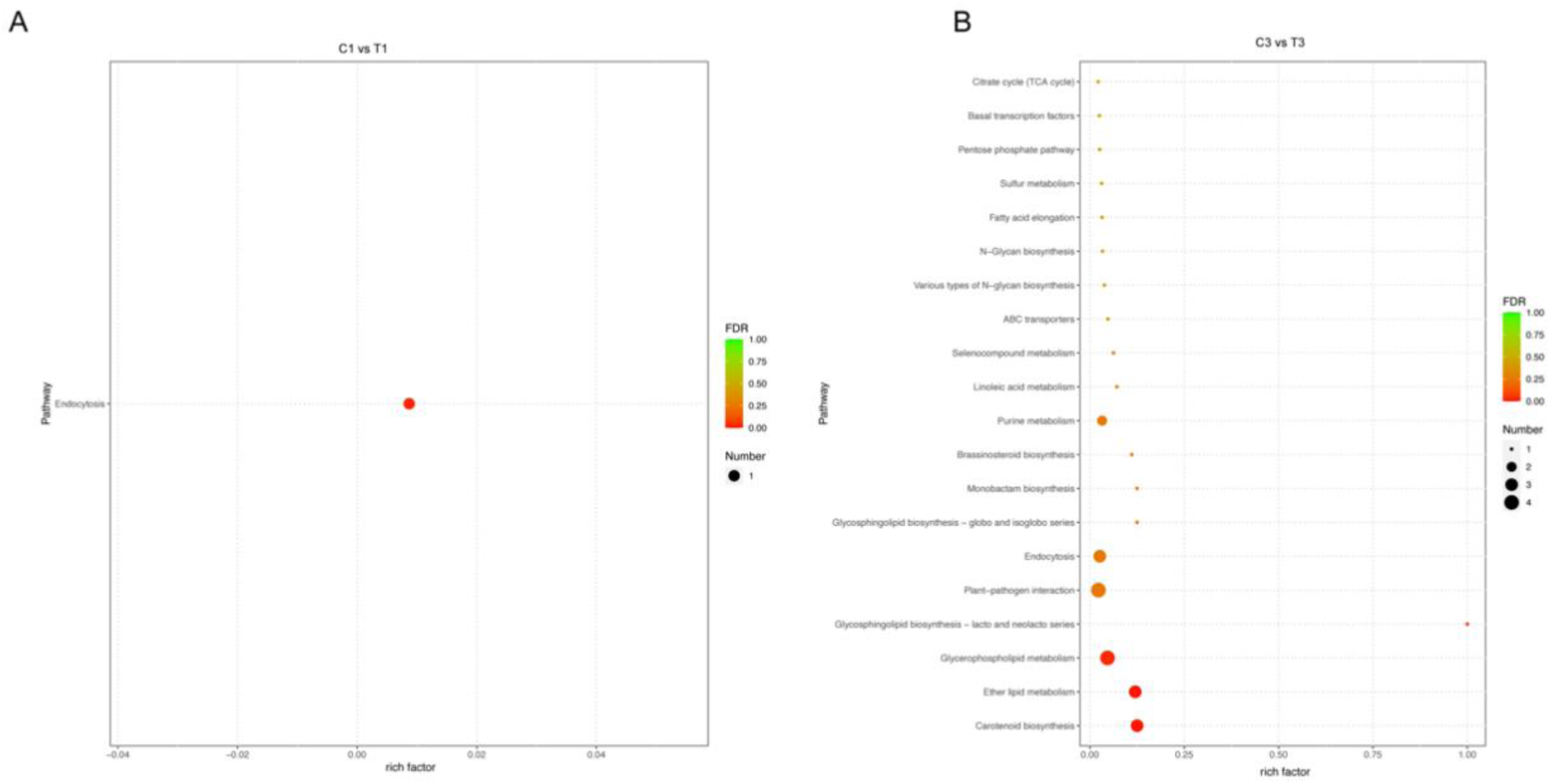
KEGG enrichment analysis of target genes of differentially expressed miRNAs post *Bacillus velezensis* 9912 treatment. (A) KEGG enrichment analysis of target genes of differentially expressed miRNAs in C1 vs T1. (B) KEGG enrichment analysis of target genes of differentially expressed miRNAs in C3 vs T3. KEGG enrichment analysis of the target genes of differentially expressed miRNAs was determined by the number of target genes included in various KEGG pathways at different hierarchical levels. The level of enrichment was measured based on the Rich factor, FDR value, and the number of miRNA target genes enriched in a specific KEGG pathway. The most significantly enriched pathways twenty KEGG pathways with the lowest FDR values were presented in KEGG enrichment results. C1, T1, C3, and T3 represent libraries constructed from rice seedlings treated with ddH_2_O (CK) and *B. velezensis* 9912 and collected at 1 and 3 days, respectively.

**Table S1.**
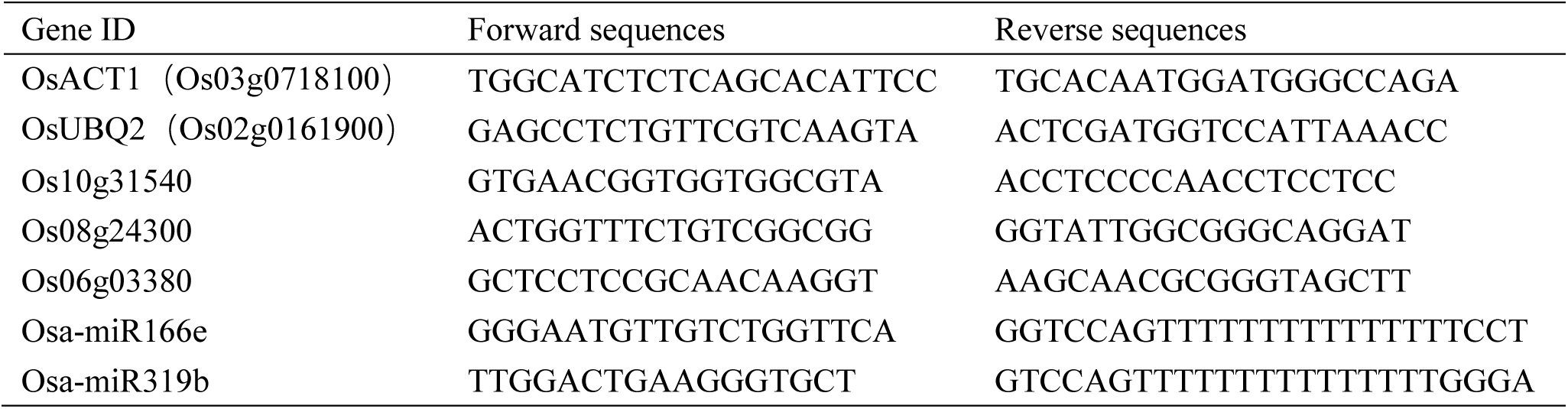
Primers used for qRT-PCR in this study.

**Table S2.** Differential expression gene in C1 *vs* T1 and C3 *vs* T3.

| Gene ID | Annotation function | Log<sub>2</sub>(FC) |  |
| --- | --- | --- | --- |
|  |  | C1 vs T1 | C3 vs T3 |
| LOC_Os01g48446 | NAC domain-containing protein 68 | -1.07 | -1.24 |
| LOC_Os09g35010 | Dehydration-responsive element-binding protein 1B | -1.75 | -2.00 |
| LOC_Os09g35020 | Dehydration-responsive element-binding protein 1H | -2.06 | -4.53 |
| LOC_Os04g55100 | Uncharacterized | -1.18 | -1.77 |
| LOC_Os02g51440 | Laccase-9 | 1.61 | 3.44 |
| LOC_Os04g55740 | Peroxidase 4 | 1.87 | 2.53 |
| LOC_Os07g44550 | Peroxidase 2 | 1.78 | 3.96 |
| LOC_Os03g21960 | Alanine--glyoxylate aminotransferase 2 homolog 3, mitochondrial | 1.40 | 1.65 |
| LOC_Os04g03579 | *L*-type lectin-domain containing receptor kinase IX.1 | 2.28 | 3.45 |
| LOC_Os04g11040 | Hypothetical protein OsJ_13938 | 2.23 | 3.63 |
| LOC_Os05g08620 | Small polypeptide DEVIL 10 | 1.81 | 1.56 |
| LOC_Os09g19350 | Probable LRR receptor-like serine/threonine-protein kinase At1g51880 | 1.40 | 2.37 |
| LOC_Os01g48660 | Uncharacterized | 1.35 | 1.82 |
| LOC_Os03g30580 | Uncharacterized | 1.02 | -1.77 |
| LOC_Os06g33450 | Spch_Arath Transcription factor Speechless OS | 1.17 | -2.62 |
| LOC_Os01g50170 | NEP1_NEPGR Aspartic proteinase nepenthesin-1 OS | -1.40 | 1.58 |
| LOC_Os01g64440 | Uncharacterized | -1.39 | 2.40 |
| LOC_Os01g68300 | Uncharacterized | -1.58 | 2.00 |
| LOC_Os01g70490 | HAK5_ORYSJ Potassium transporter 5 OS | -1.11 | 1.16 |
| LOC_Os01g73250 | ASR2_SOLLC Absciscic stress-ripening protein 2 OS | -1.44 | 3.75 |
| LOC_Os02g02230 | CP51_SORBI Obtusifolius 14-α demethylase OS | / | 4.10 |
| LOC_Os02g26430 | WRK51_ORYSJ WRKY transcription factor WRKY51 OS | -1.74 | 1.40 |
| LOC_Os02g30110 | CYQ32_CATRO Cytochrome P450 81Q32 OS | -1.24 | 1.19 |
| LOC_Os02g36110 | C76M7_ORYSJ Ent-cassadiene C11-α-hydroxylase 1 OS | -1.70 | 2.72 |
| LOC_Os02g36140 | KSL7_ORYSJ Ent-cassa-12, 15-diene synthase OS | -1.33 | 2.21 |
| LOC_Os02g36190 | C71Z7_ORYSJ Ent-cassadiene hydroxylase OS | -1.85 | 3.36 |
| LOC_Os02g36280 | C76M6_ORYSJ Oryzalexin E synthase OS | -1.63 | 3.28 |
| LOC_Os03g45860 | SAU71_ARATH Auxin-responsive protein SAUR71 OS | -1.38 | 3.93 |
| LOC_Os04g08550 | DMAS1_MAIZE Deoxymugineic acid synthase 1 OS | -1.10 | 2.17 |
| LOC_Os04g10160 | C99A2_ORYSJ Cytochrome P450 99A2 OS | -1.15 | 1.50 |
| LOC_Os04g22120 | LRK91_ARATH *L*-type lectin-domain containing receptor kinase IX.1 OS | -2.74 | 4.19 |
| LOC_Os04g30390 | Uncharacterized | -1.80 | 1.29 |
| LOC_Os04g31760 | Hypothetical protein DAI22_04g105200 | -1.94 | 2.15 |
| LOC_Os05g24780 | CML21_ORYSJ Probable calcium-binding protein CML21 OS | -1.30 | 1.15 |
| LOC_Os07g07290 | Hypothetical protein | -2.65 | 2.19 |
| LOC_Os07g07300 | Uncharacterized | -1.01 | 2.07 |
| LOC_Os07g13480 | Hypothetical protein OsI_25468 | -1.32 | 1.69 |
| LOC_Os07g23410 | DES2_SORBI Fatty acid desaturase DES2 OS | -1.18 | 3.36 |
| LOC_Os07g23430 | DES2_SORBI Fatty acid desaturase DES2 OS | -1.35 | 3.33 |
| LOC_Os08g29910 | COQ2_ARATH 4-hydroxybenzoate polyprenyltransferase, mitochondrial OS | -2.75 | 1.42 |
| LOC_Os09g17560 | ASMT1_ORYSJ Acetylserotonin *O*-methyltransferase 1 OS | -1.83 | 4.93 |
| LOC_Os09g18260 | Y1518_ARATH Probable LRR receptor-like serine/threonine-protein kinase At1g51860 OS | -1.61 | 4.01 |
| LOC_Os10g04570 | RGA2_SOLBU Disease resistance protein RGA2 OS | -4.69 | 2.77 |
| LOC_Os11g06550 | Hypothetical protein LOC_Os11g06550 | -1.47 | 4.04 |
| LOC_Os11g31540 | BAK1_ORYSJ LRR receptor kinase BAK1 OS | -1.28 | 2.13 |
| LOC_Os11g37210 | Neurogenic locus notch homolog protein 1 | -1.07 | 1.18 |
| LOC_Os12g02400 | WRK54_ARATH Probable WRKY transcription factor 54 OS | -4.35 | 3.96 |
| LOC_Os12g11620 | Hypothetical protein LOC_Os12g11620 | -1.44 | 1.29 |
| LOC_Os12g11980 | Hypothetical protein LOC_Os12g11980 | -1.44 | 1.20 |
| LOC_Os12g12120 | RLP35_ARATH Receptor-like protein 35 OS | -1.20 | 1.99 |
| LOC_Os12g42280 | NCED5_ORYSJ 9-cis-epoxycarotenoid dioxygenase NCED5, chloroplastic OS | -1.77 | 1.83 |

Note: Log<sub>2</sub>(FC) indicates log<sub>2</sub>(FoldChange). C1, T1, C3, and T3 represent libraries constructed from rice seedlings treated with ddH<sub>2</sub>O (CK) and *B. velezensis* 9912 and collected at 1 and 3 days, respectively.

**Table S3.** The differentially expressed miRNAs in rice seedlings after *B. velezensis* 9912 treatment.

| Comparison set | Gene ID | FoldChange | Log<sub>2</sub>(FC) | P-value | Regulation |
| --- | --- | --- | --- | --- | --- |
| C1 vs T1 | osa-miR5799 | 21.7 | 4.4 | 0.008 | Up |
|  | osa-miR821c | Inf | Inf | 0.025 | Up |
| C3 vs T3 | osa-miR1432-3p | 4.0 | 2.0 | 0.031 | Up |
|  | osa-miR1440b | 24.4 | 4.6 | 0.043 | Up |
|  | osa-miR2873a | 7.9 | 3.0 | 7.76E-07 | Up |
|  | osa-miR2907d | 35.6 | 5.2 | 0.022 | Up |
|  | osa-miR395y | 2.6 | 1.4 | 5.51E-05 | Up |
|  | osa-miR397b | 4.8 | 2.3 | 0.003 | Up |
|  | osa-miR5155 | 11.1 | 3.5 | 0.033 | Up |
|  | osa-miR6251 | 2.3 | 1.2 | 0.001 | Up |
|  | osa-miR827 | 4.0 | 2.0 | 0.0006 | Up |
|  | osa-miR166e-5p | 0.3 | -1.6 | 0.0002 | Down |
|  | osa-miR166h-5p | 0.4 | -1.5 | 0.0002 | Down |
|  | osa-miR166l-5p | 0.4 | -1.3 | 0.0024 | Down |
|  | osa-miR171c-5p | 0.5 | -1.0 | 0.018 | Down |
|  | osa-miR171e-5p | 0.5 | -1.1 | 0.0081 | Down |
|  | osa-miR171f-5p | 0.2 | -2.6 | 0.0027 | Down |
|  | osa-miR171h | 0.2 | -2.5 | 0.031 | Down |
|  | osa-miR1859 | 0.3 | -1.5 | 0.0041 | Down |
|  | osa-miR1861l | 0.2 | -2.7 | 0.0054 | Down |
|  | osa-miR2867-3p | 0 | / | 0.0044 | Down |
|  | osa-miR319a-3p | 0.2 | -2.0 | 2.23E-06 | Down |
|  | osa-miR319b | 0.2 | -2.3 | 4.95E-15 | Down |
|  | osa-miR396b-3p | 0.2 | -2.2 | 0.047 | Down |
|  | osa-miR396b-5p | 0.4 | -1.2 | 2.52E-06 | Down |
|  | osa-miR3981-5p | 0 | / | 0.031 | Down |
|  | osa-miR437 | 0.1 | -2.8 | 0.044 | Down |
|  | osa-miR5081 | 0.2 | -2.2 | 0.035 | Down |
|  | osa-miR529a | 0.4 | -1.2 | 8.80E-05 | Down |
|  | osa-miR6249b | 0.2 | -2.5 | 0.024 | Down |

Note: DESeq software was used to analyze the differentially expressed miRNA. miRNAs with $|\log_2\text{FoldChange}| > 1$ and significance $p\text{-value} < 0.05$ were assigned as considered as differentially expressed. C1, T1, C3, and T3 represent libraries constructed from rice seedlings treated with ddH<sub>2</sub>O (CK) and *B. velezensis* 9912 and collected at days 1 and 3, respectively.

**Table S4.**
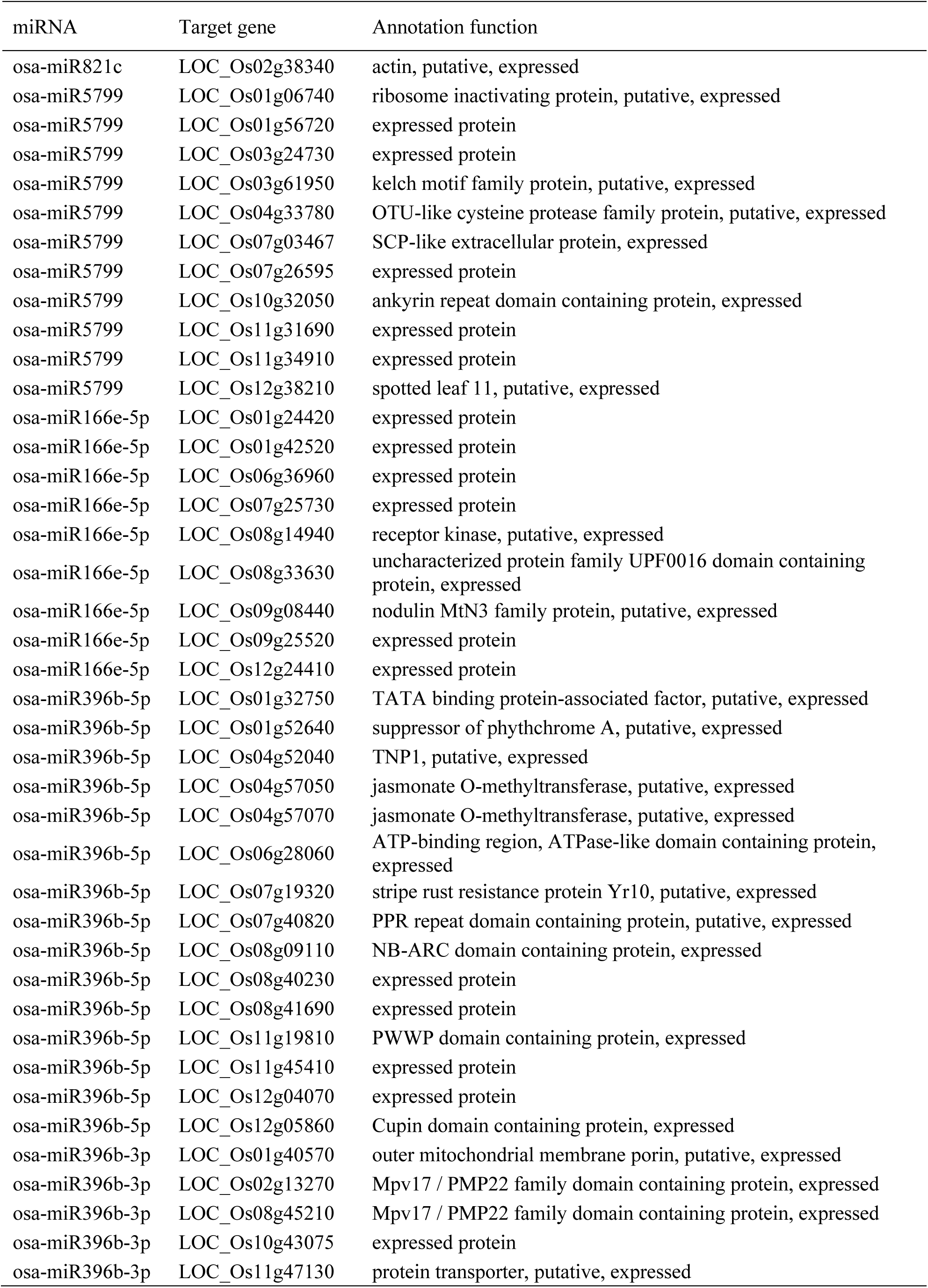
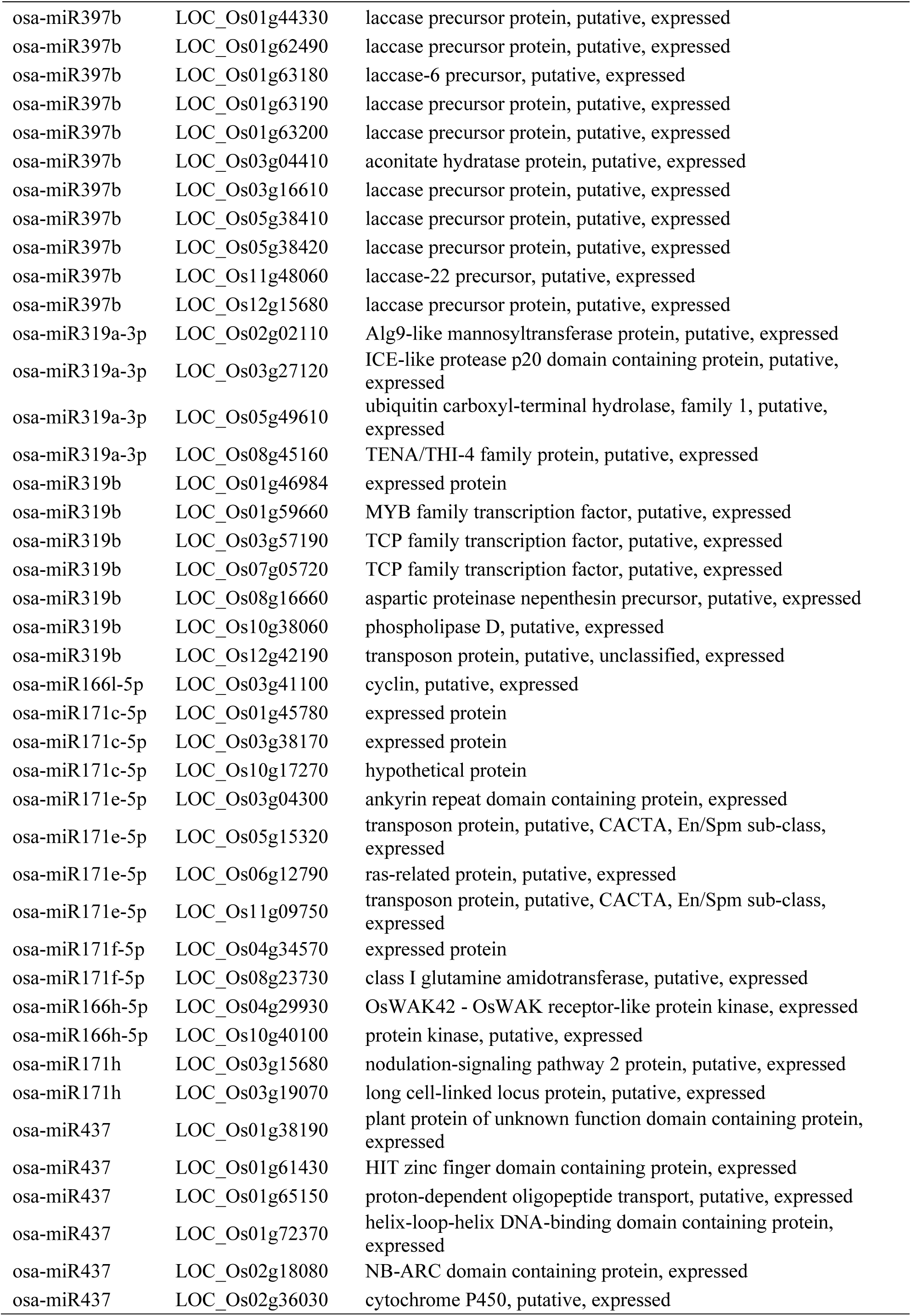
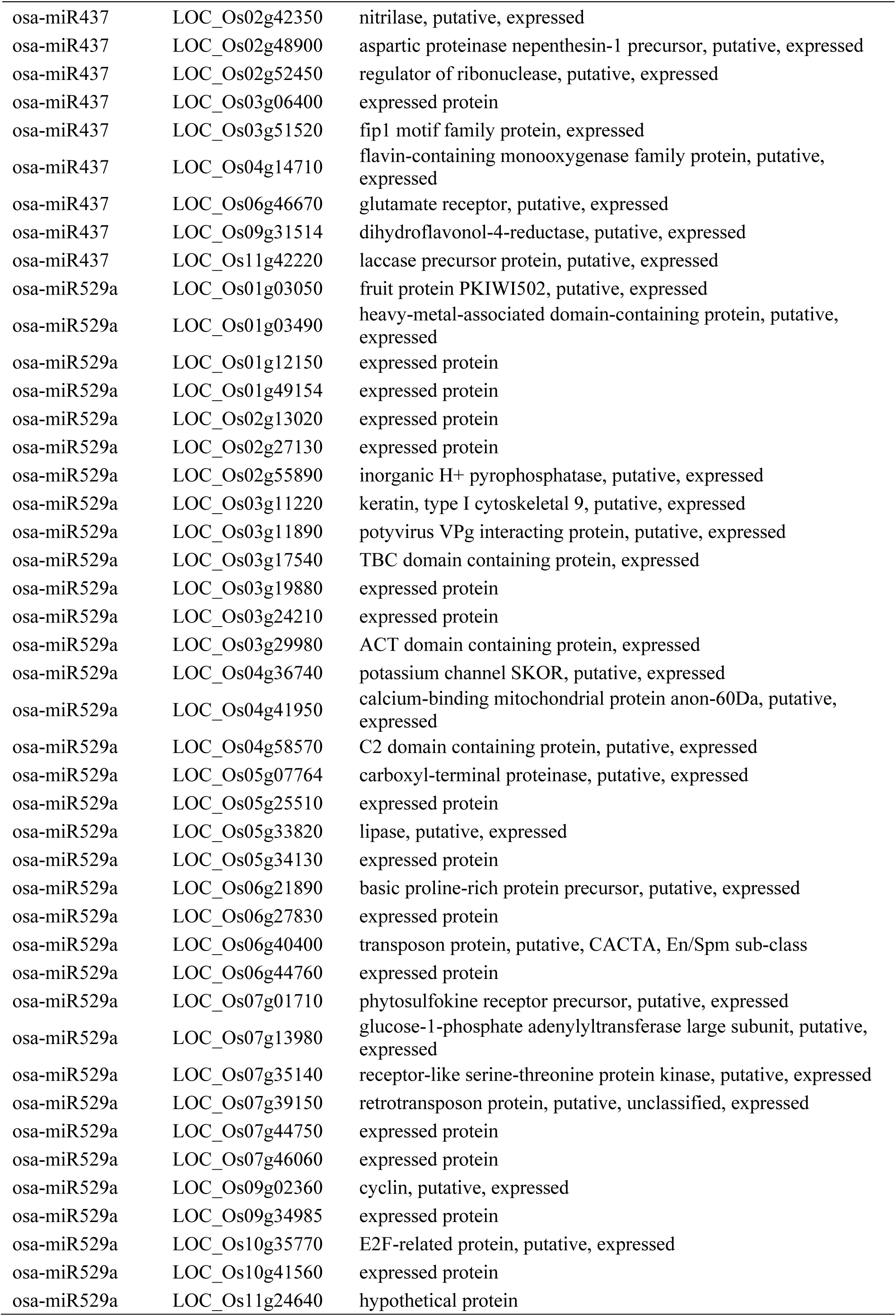
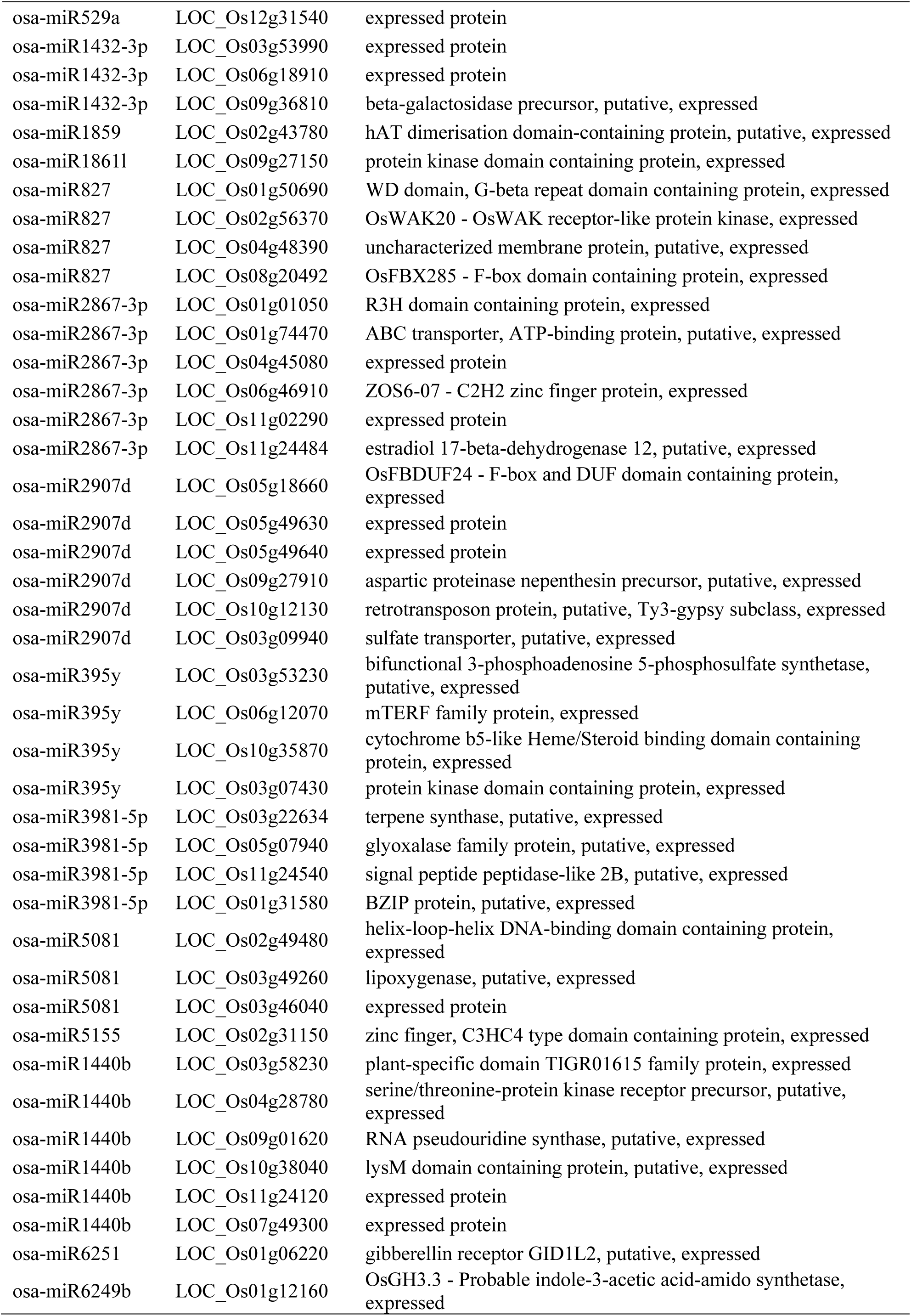
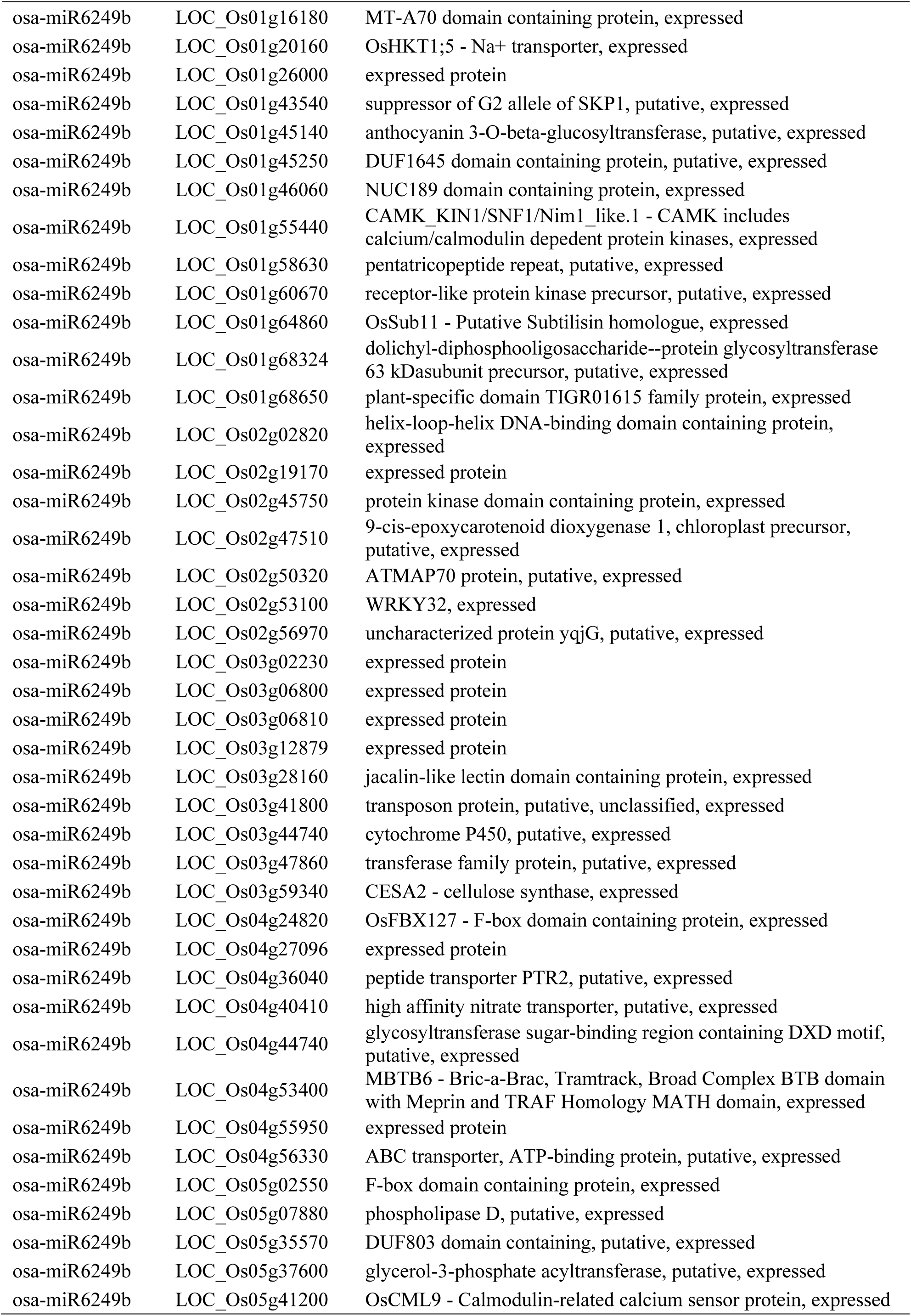
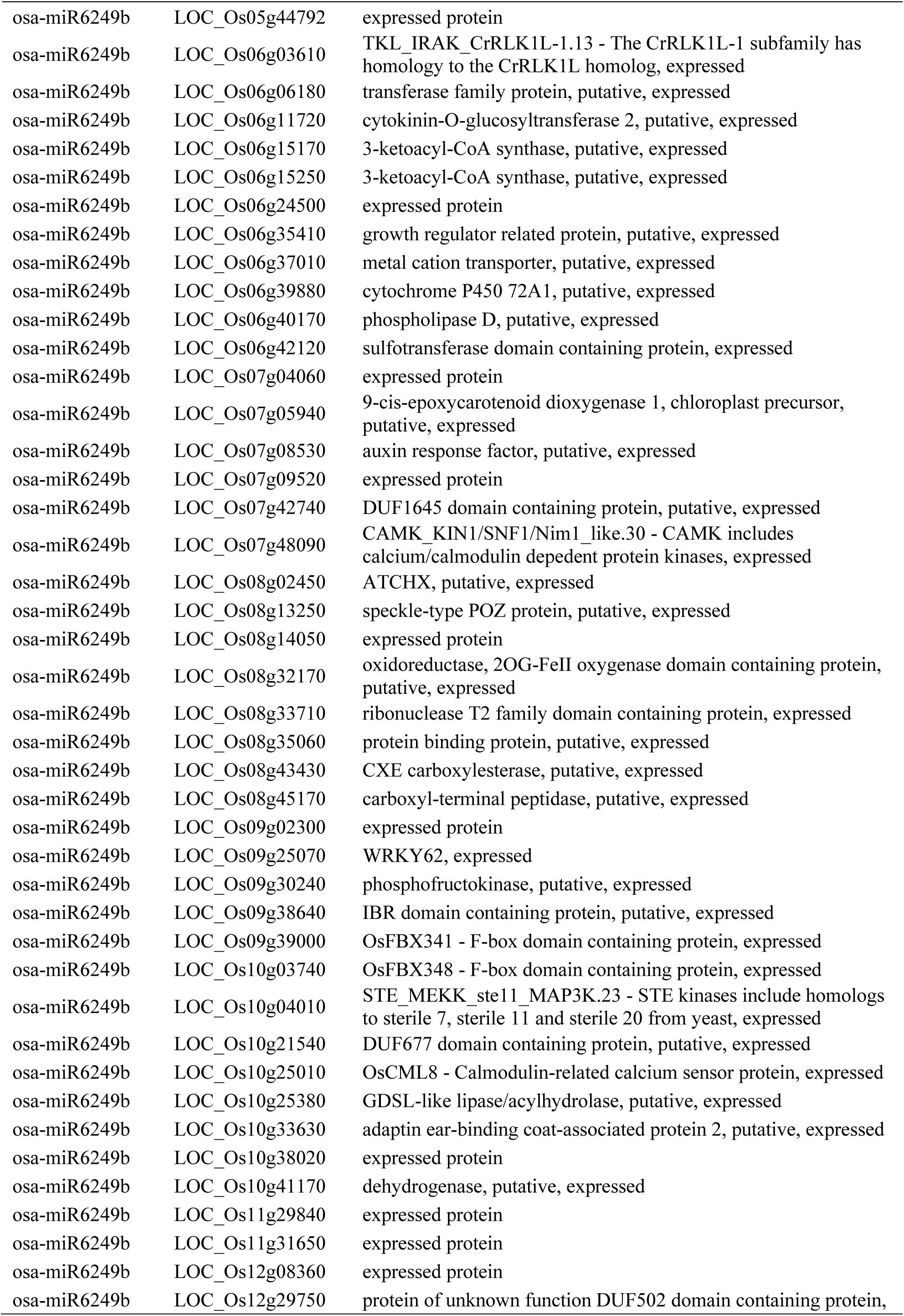
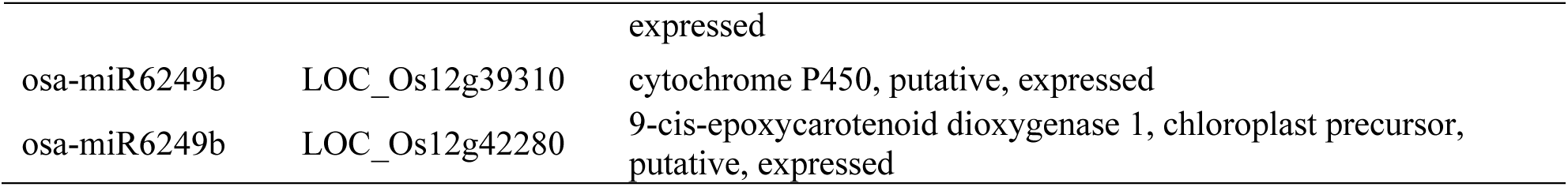
Target genes of differentially expressed miRNAs.

| miRNA | Target gene | Annotation function |
| --- | --- | --- |
| osa-miR821c | LOC_Os02g38340 | actin, putative, expressed |
| osa-miR5799 | LOC_Os01g06740 | ribosome inactivating protein, putative, expressed |
| osa-miR5799 | LOC_Os01g56720 | expressed protein |
| osa-miR5799 | LOC_Os03g24730 | expressed protein |
| osa-miR5799 | LOC_Os03g61950 | kelch motif family protein, putative, expressed |
| osa-miR5799 | LOC_Os04g33780 | OTU-like cysteine protease family protein, putative, expressed |
| osa-miR5799 | LOC_Os07g03467 | SCP-like extracellular protein, expressed |
| osa-miR5799 | LOC_Os07g26595 | expressed protein |
| osa-miR5799 | LOC_Os10g32050 | ankyrin repeat domain containing protein, expressed |
| osa-miR5799 | LOC_Os11g31690 | expressed protein |
| osa-miR5799 | LOC_Os11g34910 | expressed protein |
| osa-miR5799 | LOC_Os12g38210 | spotted leaf 11, putative, expressed |
| osa-miR166e-5p | LOC_Os01g24420 | expressed protein |
| osa-miR166e-5p | LOC_Os01g42520 | expressed protein |
| osa-miR166e-5p | LOC_Os06g36960 | expressed protein |
| osa-miR166e-5p | LOC_Os07g25730 | expressed protein |
| osa-miR166e-5p | LOC_Os08g14940 | receptor kinase, putative, expressed |
| osa-miR166e-5p | LOC_Os08g33630 | uncharacterized protein family UPF0016 domain containing protein, expressed |
| osa-miR166e-5p | LOC_Os09g08440 | nodulin MtN3 family protein, putative, expressed |
| osa-miR166e-5p | LOC_Os09g25520 | expressed protein |
| osa-miR166e-5p | LOC_Os12g24410 | expressed protein |
| osa-miR396b-5p | LOC_Os01g32750 | TATA binding protein-associated factor, putative, expressed |
| osa-miR396b-5p | LOC_Os01g52640 | suppressor of phythochrome A, putative, expressed |
| osa-miR396b-5p | LOC_Os04g52040 | TNP1, putative, expressed |
| osa-miR396b-5p | LOC_Os04g57050 | jasmonate O-methyltransferase, putative, expressed |
| osa-miR396b-5p | LOC_Os04g57070 | jasmonate O-methyltransferase, putative, expressed |
| osa-miR396b-5p | LOC_Os06g28060 | ATP-binding region, ATPase-like domain containing protein, expressed |
| osa-miR396b-5p | LOC_Os07g19320 | stripe rust resistance protein Yr10, putative, expressed |
| osa-miR396b-5p | LOC_Os07g40820 | PPR repeat domain containing protein, putative, expressed |
| osa-miR396b-5p | LOC_Os08g09110 | NB-ARC domain containing protein, expressed |
| osa-miR396b-5p | LOC_Os08g40230 | expressed protein |
| osa-miR396b-5p | LOC_Os08g41690 | expressed protein |
| osa-miR396b-5p | LOC_Os11g19810 | PWWP domain containing protein, expressed |
| osa-miR396b-5p | LOC_Os11g45410 | expressed protein |
| osa-miR396b-5p | LOC_Os12g04070 | expressed protein |
| osa-miR396b-5p | LOC_Os12g05860 | Cupin domain containing protein, expressed |
| osa-miR396b-3p | LOC_Os01g40570 | outer mitochondrial membrane porin, putative, expressed |
| osa-miR396b-3p | LOC_Os02g13270 | Mpv17 / PMP22 family domain containing protein, expressed |
| osa-miR396b-3p | LOC_Os08g45210 | Mpv17 / PMP22 family domain containing protein, expressed |
| osa-miR396b-3p | LOC_Os10g43075 | expressed protein |
| osa-miR396b-3p | LOC_Os11g47130 | protein transporter, putative, expressed |
| osa-miR397b | LOC_Os01g44330 | laccase precursor protein, putative, expressed |
| osa-miR397b | LOC_Os01g62490 | laccase precursor protein, putative, expressed |
| osa-miR397b | LOC_Os01g63180 | laccase-6 precursor, putative, expressed |
| osa-miR397b | LOC_Os01g63190 | laccase precursor protein, putative, expressed |
| osa-miR397b | LOC_Os01g63200 | laccase precursor protein, putative, expressed |
| osa-miR397b | LOC_Os03g04410 | aconitate hydratase protein, putative, expressed |
| osa-miR397b | LOC_Os03g16610 | laccase precursor protein, putative, expressed |
| osa-miR397b | LOC_Os05g38410 | laccase precursor protein, putative, expressed |
| osa-miR397b | LOC_Os05g38420 | laccase precursor protein, putative, expressed |
| osa-miR397b | LOC_Os11g48060 | laccase-22 precursor, putative, expressed |
| osa-miR397b | LOC_Os12g15680 | laccase precursor protein, putative, expressed |
| osa-miR319a-3p | LOC_Os02g02110 | Alg9-like mannosyltransferase protein, putative, expressed |
| osa-miR319a-3p | LOC_Os03g27120 | ICE-like protease p20 domain containing protein, putative, expressed |
| osa-miR319a-3p | LOC_Os05g49610 | ubiquitin carboxyl-terminal hydrolase, family 1, putative, expressed |
| osa-miR319a-3p | LOC_Os08g45160 | TENA/THI-4 family protein, putative, expressed |
| osa-miR319b | LOC_Os01g46984 | expressed protein |
| osa-miR319b | LOC_Os01g59660 | MYB family transcription factor, putative, expressed |
| osa-miR319b | LOC_Os03g57190 | TCP family transcription factor, putative, expressed |
| osa-miR319b | LOC_Os07g05720 | TCP family transcription factor, putative, expressed |
| osa-miR319b | LOC_Os08g16660 | aspartic proteinase nepenthesin precursor, putative, expressed |
| osa-miR319b | LOC_Os10g38060 | phospholipase D, putative, expressed |
| osa-miR319b | LOC_Os12g42190 | transposon protein, putative, unclassified, expressed |
| osa-miR166l-5p | LOC_Os03g41100 | cyclin, putative, expressed |
| osa-miR171c-5p | LOC_Os01g45780 | expressed protein |
| osa-miR171c-5p | LOC_Os03g38170 | expressed protein |
| osa-miR171c-5p | LOC_Os10g17270 | hypothetical protein |
| osa-miR171e-5p | LOC_Os03g04300 | ankyrin repeat domain containing protein, expressed |
| osa-miR171e-5p | LOC_Os05g15320 | transposon protein, putative, CACTA, En/Spm sub-class, expressed |
| osa-miR171e-5p | LOC_Os06g12790 | ras-related protein, putative, expressed |
| osa-miR171e-5p | LOC_Os11g09750 | transposon protein, putative, CACTA, En/Spm sub-class, expressed |
| osa-miR171f-5p | LOC_Os04g34570 | expressed protein |
| osa-miR171f-5p | LOC_Os08g23730 | class I glutamine amidotransferase, putative, expressed |
| osa-miR166h-5p | LOC_Os04g29930 | OsWAK42 - OsWAK receptor-like protein kinase, expressed |
| osa-miR166h-5p | LOC_Os10g40100 | protein kinase, putative, expressed |
| osa-miR171h | LOC_Os03g15680 | nodulation-signaling pathway 2 protein, putative, expressed |
| osa-miR171h | LOC_Os03g19070 | long cell-linked locus protein, putative, expressed |
| osa-miR437 | LOC_Os01g38190 | plant protein of unknown function domain containing protein, expressed |
| osa-miR437 | LOC_Os01g61430 | HIT zinc finger domain containing protein, expressed |
| osa-miR437 | LOC_Os01g65150 | proton-dependent oligopeptide transport, putative, expressed |
| osa-miR437 | LOC_Os01g72370 | helix-loop-helix DNA-binding domain containing protein, expressed |
| osa-miR437 | LOC_Os02g18080 | NB-ARC domain containing protein, expressed |
| osa-miR437 | LOC_Os02g36030 | cytochrome P450, putative, expressed |
| osa-miR437 | LOC_Os02g42350 | nitrilase, putative, expressed |
| osa-miR437 | LOC_Os02g48900 | aspartic proteinase nepenthesin-1 precursor, putative, expressed |
| osa-miR437 | LOC_Os02g52450 | regulator of ribonuclease, putative, expressed |
| osa-miR437 | LOC_Os03g06400 | expressed protein |
| osa-miR437 | LOC_Os03g51520 | fip1 motif family protein, expressed |
| osa-miR437 | LOC_Os04g14710 | flavin-containing monooxygenase family protein, putative, expressed |
| osa-miR437 | LOC_Os06g46670 | glutamate receptor, putative, expressed |
| osa-miR437 | LOC_Os09g31514 | dihydroflavonol-4-reductase, putative, expressed |
| osa-miR437 | LOC_Os11g42220 | laccase precursor protein, putative, expressed |
| osa-miR529a | LOC_Os01g03050 | fruit protein PKIWI502, putative, expressed |
| osa-miR529a | LOC_Os01g03490 | heavy-metal-associated domain-containing protein, putative, expressed |
| osa-miR529a | LOC_Os01g12150 | expressed protein |
| osa-miR529a | LOC_Os01g49154 | expressed protein |
| osa-miR529a | LOC_Os02g13020 | expressed protein |
| osa-miR529a | LOC_Os02g27130 | expressed protein |
| osa-miR529a | LOC_Os02g55890 | inorganic H<sup>+</sup> pyrophosphatase, putative, expressed |
| osa-miR529a | LOC_Os03g11220 | keratin, type I cytoskeletal 9, putative, expressed |
| osa-miR529a | LOC_Os03g11890 | potyvirus VPg interacting protein, putative, expressed |
| osa-miR529a | LOC_Os03g17540 | TBC domain containing protein, expressed |
| osa-miR529a | LOC_Os03g19880 | expressed protein |
| osa-miR529a | LOC_Os03g24210 | expressed protein |
| osa-miR529a | LOC_Os03g29980 | ACT domain containing protein, expressed |
| osa-miR529a | LOC_Os04g36740 | potassium channel SKOR, putative, expressed |
| osa-miR529a | LOC_Os04g41950 | calcium-binding mitochondrial protein anon-60Da, putative, expressed |
| osa-miR529a | LOC_Os04g58570 | C2 domain containing protein, putative, expressed |
| osa-miR529a | LOC_Os05g07764 | carboxyl-terminal proteinase, putative, expressed |
| osa-miR529a | LOC_Os05g25510 | expressed protein |
| osa-miR529a | LOC_Os05g33820 | lipase, putative, expressed |
| osa-miR529a | LOC_Os05g34130 | expressed protein |
| osa-miR529a | LOC_Os06g21890 | basic proline-rich protein precursor, putative, expressed |
| osa-miR529a | LOC_Os06g27830 | expressed protein |
| osa-miR529a | LOC_Os06g40400 | transposon protein, putative, CACTA, En/Spm sub-class |
| osa-miR529a | LOC_Os06g44760 | expressed protein |
| osa-miR529a | LOC_Os07g01710 | phytosulfokine receptor precursor, putative, expressed |
| osa-miR529a | LOC_Os07g13980 | glucose-1-phosphate adenylyltransferase large subunit, putative, expressed |
| osa-miR529a | LOC_Os07g35140 | receptor-like serine-threonine protein kinase, putative, expressed |
| osa-miR529a | LOC_Os07g39150 | retrotransposon protein, putative, unclassified, expressed |
| osa-miR529a | LOC_Os07g44750 | expressed protein |
| osa-miR529a | LOC_Os07g46060 | expressed protein |
| osa-miR529a | LOC_Os09g02360 | cyclin, putative, expressed |
| osa-miR529a | LOC_Os09g34985 | expressed protein |
| osa-miR529a | LOC_Os10g35770 | E2F-related protein, putative, expressed |
| osa-miR529a | LOC_Os10g41560 | expressed protein |
| osa-miR529a | LOC_Os11g24640 | hypothetical protein |
| osa-miR529a | LOC_Os12g31540 | expressed protein |
| osa-miR1432-3p | LOC_Os03g53990 | expressed protein |
| osa-miR1432-3p | LOC_Os06g18910 | expressed protein |
| osa-miR1432-3p | LOC_Os09g36810 | beta-galactosidase precursor, putative, expressed |
| osa-miR1859 | LOC_Os02g43780 | hAT dimerisation domain-containing protein, putative, expressed |
| osa-miR1861l | LOC_Os09g27150 | protein kinase domain containing protein, expressed |
| osa-miR827 | LOC_Os01g50690 | WD domain, G-beta repeat domain containing protein, expressed |
| osa-miR827 | LOC_Os02g56370 | OsWAK20 - OsWAK receptor-like protein kinase, expressed |
| osa-miR827 | LOC_Os04g48390 | uncharacterized membrane protein, putative, expressed |
| osa-miR827 | LOC_Os08g20492 | OsFBX285 - F-box domain containing protein, expressed |
| osa-miR2867-3p | LOC_Os01g01050 | R3H domain containing protein, expressed |
| osa-miR2867-3p | LOC_Os01g74470 | ABC transporter, ATP-binding protein, putative, expressed |
| osa-miR2867-3p | LOC_Os04g45080 | expressed protein |
| osa-miR2867-3p | LOC_Os06g46910 | ZOS6-07 - C2H2 zinc finger protein, expressed |
| osa-miR2867-3p | LOC_Os11g02290 | expressed protein |
| osa-miR2867-3p | LOC_Os11g24484 | estradiol 17-beta-dehydrogenase 12, putative, expressed |
| osa-miR2907d | LOC_Os05g18660 | OsFBDUF24 - F-box and DUF domain containing protein, expressed |
| osa-miR2907d | LOC_Os05g49630 | expressed protein |
| osa-miR2907d | LOC_Os05g49640 | expressed protein |
| osa-miR2907d | LOC_Os09g27910 | aspartic proteinase nepenthesin precursor, putative, expressed |
| osa-miR2907d | LOC_Os10g12130 | retrotransposon protein, putative, Ty3-gypsy subclass, expressed |
| osa-miR2907d | LOC_Os03g09940 | sulfate transporter, putative, expressed |
| osa-miR395y | LOC_Os03g53230 | bifunctional 3-phosphoadenosine 5-phosphosulfate synthetase, putative, expressed |
| osa-miR395y | LOC_Os06g12070 | mTERF family protein, expressed |
| osa-miR395y | LOC_Os10g35870 | cytochrome b5-like Heme/Steroid binding domain containing protein, expressed |
| osa-miR395y | LOC_Os03g07430 | protein kinase domain containing protein, expressed |
| osa-miR3981-5p | LOC_Os03g22634 | terpene synthase, putative, expressed |
| osa-miR3981-5p | LOC_Os05g07940 | glyoxalase family protein, putative, expressed |
| osa-miR3981-5p | LOC_Os11g24540 | signal peptide peptidase-like 2B, putative, expressed |
| osa-miR3981-5p | LOC_Os01g31580 | BZIP protein, putative, expressed |
| osa-miR5081 | LOC_Os02g49480 | helix-loop-helix DNA-binding domain containing protein, expressed |
| osa-miR5081 | LOC_Os03g49260 | lipoxygenase, putative, expressed |
| osa-miR5081 | LOC_Os03g46040 | expressed protein |
| osa-miR5155 | LOC_Os02g31150 | zinc finger, C3HC4 type domain containing protein, expressed |
| osa-miR1440b | LOC_Os03g58230 | plant-specific domain TIGR01615 family protein, expressed |
| osa-miR1440b | LOC_Os04g28780 | serine/threonine-protein kinase receptor precursor, putative, expressed |
| osa-miR1440b | LOC_Os09g01620 | RNA pseudouridine synthase, putative, expressed |
| osa-miR1440b | LOC_Os10g38040 | lysM domain containing protein, putative, expressed |
| osa-miR1440b | LOC_Os11g24120 | expressed protein |
| osa-miR1440b | LOC_Os07g49300 | expressed protein |
| osa-miR6251 | LOC_Os01g06220 | gibberellin receptor GID1L2, putative, expressed |
| osa-miR6249b | LOC_Os01g12160 | OsGH3.3 - Probable indole-3-acetic acid-amido synthetase, expressed |
| osa-miR6249b | LOC_Os01g16180 | MT-A70 domain containing protein, expressed |
| osa-miR6249b | LOC_Os01g20160 | OsHKT1;5 - Na<sup>+</sup> transporter, expressed |
| osa-miR6249b | LOC_Os01g26000 | expressed protein |
| osa-miR6249b | LOC_Os01g43540 | suppressor of G2 allele of SKP1, putative, expressed |
| osa-miR6249b | LOC_Os01g45140 | anthocyanin 3-O-beta-glucosyltransferase, putative, expressed |
| osa-miR6249b | LOC_Os01g45250 | DUF1645 domain containing protein, putative, expressed |
| osa-miR6249b | LOC_Os01g46060 | NUC189 domain containing protein, expressed |
| osa-miR6249b | LOC_Os01g55440 | CAMK_KIN1/SNF1/Nim1_like.1 - CAMK includes calcium/calmodulin dependent protein kinases, expressed |
| osa-miR6249b | LOC_Os01g58630 | pentatricopeptide repeat, putative, expressed |
| osa-miR6249b | LOC_Os01g60670 | receptor-like protein kinase precursor, putative, expressed |
| osa-miR6249b | LOC_Os01g64860 | OsSub11 - Putative Subtilisin homologue, expressed |
| osa-miR6249b | LOC_Os01g68324 | dolichyl-diphosphooligosaccharide--protein glycosyltransferase 63 kDa subunit precursor, putative, expressed |
| osa-miR6249b | LOC_Os01g68650 | plant-specific domain TIGR01615 family protein, expressed |
| osa-miR6249b | LOC_Os02g02820 | helix-loop-helix DNA-binding domain containing protein, expressed |
| osa-miR6249b | LOC_Os02g19170 | expressed protein |
| osa-miR6249b | LOC_Os02g45750 | protein kinase domain containing protein, expressed |
| osa-miR6249b | LOC_Os02g47510 | 9-cis-epoxycarotenoid dioxygenase 1, chloroplast precursor, putative, expressed |
| osa-miR6249b | LOC_Os02g50320 | ATMAP70 protein, putative, expressed |
| osa-miR6249b | LOC_Os02g53100 | WRKY32, expressed |
| osa-miR6249b | LOC_Os02g56970 | uncharacterized protein yqjG, putative, expressed |
| osa-miR6249b | LOC_Os03g02230 | expressed protein |
| osa-miR6249b | LOC_Os03g06800 | expressed protein |
| osa-miR6249b | LOC_Os03g06810 | expressed protein |
| osa-miR6249b | LOC_Os03g12879 | expressed protein |
| osa-miR6249b | LOC_Os03g28160 | jacalin-like lectin domain containing protein, expressed |
| osa-miR6249b | LOC_Os03g41800 | transposon protein, putative, unclassified, expressed |
| osa-miR6249b | LOC_Os03g44740 | cytochrome P450, putative, expressed |
| osa-miR6249b | LOC_Os03g47860 | transferase family protein, putative, expressed |
| osa-miR6249b | LOC_Os03g59340 | CESA2 - cellulose synthase, expressed |
| osa-miR6249b | LOC_Os04g24820 | OsFBX127 - F-box domain containing protein, expressed |
| osa-miR6249b | LOC_Os04g27096 | expressed protein |
| osa-miR6249b | LOC_Os04g36040 | peptide transporter PTR2, putative, expressed |
| osa-miR6249b | LOC_Os04g40410 | high affinity nitrate transporter, putative, expressed |
| osa-miR6249b | LOC_Os04g44740 | glycosyltransferase sugar-binding region containing DXD motif, putative, expressed |
| osa-miR6249b | LOC_Os04g53400 | MBTB6 - Bric-a-Brac, Tramtrack, Broad Complex BTB domain with Meprin and TRAF Homology MATH domain, expressed |
| osa-miR6249b | LOC_Os04g55950 | expressed protein |
| osa-miR6249b | LOC_Os04g56330 | ABC transporter, ATP-binding protein, putative, expressed |
| osa-miR6249b | LOC_Os05g02550 | F-box domain containing protein, expressed |
| osa-miR6249b | LOC_Os05g07880 | phospholipase D, putative, expressed |
| osa-miR6249b | LOC_Os05g35570 | DUF803 domain containing, putative, expressed |
| osa-miR6249b | LOC_Os05g37600 | glycerol-3-phosphate acyltransferase, putative, expressed |
| osa-miR6249b | LOC_Os05g41200 | OsCML9 - Calmodulin-related calcium sensor protein, expressed |
| osa-miR6249b | LOC_Os05g44792 | expressed protein |
| osa-miR6249b | LOC_Os06g03610 | TKL_IRAK_CrRLK1L-1.13 - The CrRLK1L-1 subfamily has homology to the CrRLK1L homolog, expressed |
| osa-miR6249b | LOC_Os06g06180 | transferase family protein, putative, expressed |
| osa-miR6249b | LOC_Os06g11720 | cytokinin-O-glucosyltransferase 2, putative, expressed |
| osa-miR6249b | LOC_Os06g15170 | 3-ketoacyl-CoA synthase, putative, expressed |
| osa-miR6249b | LOC_Os06g15250 | 3-ketoacyl-CoA synthase, putative, expressed |
| osa-miR6249b | LOC_Os06g24500 | expressed protein |
| osa-miR6249b | LOC_Os06g35410 | growth regulator related protein, putative, expressed |
| osa-miR6249b | LOC_Os06g37010 | metal cation transporter, putative, expressed |
| osa-miR6249b | LOC_Os06g39880 | cytochrome P450 72A1, putative, expressed |
| osa-miR6249b | LOC_Os06g40170 | phospholipase D, putative, expressed |
| osa-miR6249b | LOC_Os06g42120 | sulfotransferase domain containing protein, expressed |
| osa-miR6249b | LOC_Os07g04060 | expressed protein |
| osa-miR6249b | LOC_Os07g05940 | 9-cis-epoxycarotenoid dioxygenase 1, chloroplast precursor, putative, expressed |
| osa-miR6249b | LOC_Os07g08530 | auxin response factor, putative, expressed |
| osa-miR6249b | LOC_Os07g09520 | expressed protein |
| osa-miR6249b | LOC_Os07g42740 | DUF1645 domain containing protein, putative, expressed |
| osa-miR6249b | LOC_Os07g48090 | CAMK_KIN1/SNF1/Nim1_like.30 - CAMK includes calcium/calmodulin dependent protein kinases, expressed |
| osa-miR6249b | LOC_Os08g02450 | ATCHX, putative, expressed |
| osa-miR6249b | LOC_Os08g13250 | speckle-type POZ protein, putative, expressed |
| osa-miR6249b | LOC_Os08g14050 | expressed protein |
| osa-miR6249b | LOC_Os08g32170 | oxidoreductase, 2OG-FeII oxygenase domain containing protein, putative, expressed |
| osa-miR6249b | LOC_Os08g33710 | ribonuclease T2 family domain containing protein, expressed |
| osa-miR6249b | LOC_Os08g35060 | protein binding protein, putative, expressed |
| osa-miR6249b | LOC_Os08g43430 | CXE carboxylesterase, putative, expressed |
| osa-miR6249b | LOC_Os08g45170 | carboxyl-terminal peptidase, putative, expressed |
| osa-miR6249b | LOC_Os09g02300 | expressed protein |
| osa-miR6249b | LOC_Os09g25070 | WRKY62, expressed |
| osa-miR6249b | LOC_Os09g30240 | phosphofructokinase, putative, expressed |
| osa-miR6249b | LOC_Os09g38640 | IBR domain containing protein, putative, expressed |
| osa-miR6249b | LOC_Os09g39000 | OsFBX341 - F-box domain containing protein, expressed |
| osa-miR6249b | LOC_Os10g03740 | OsFBX348 - F-box domain containing protein, expressed |
| osa-miR6249b | LOC_Os10g04010 | STE_MEKK_ste11_MAP3K.23 - STE kinases include homologs to sterile 7, sterile 11 and sterile 20 from yeast, expressed |
| osa-miR6249b | LOC_Os10g21540 | DUF677 domain containing protein, putative, expressed |
| osa-miR6249b | LOC_Os10g25010 | OsCML8 - Calmodulin-related calcium sensor protein, expressed |
| osa-miR6249b | LOC_Os10g25380 | GDSL-like lipase/acylhydrolase, putative, expressed |
| osa-miR6249b | LOC_Os10g33630 | adaptin ear-binding coat-associated protein 2, putative, expressed |
| osa-miR6249b | LOC_Os10g38020 | expressed protein |
| osa-miR6249b | LOC_Os10g41170 | dehydrogenase, putative, expressed |
| osa-miR6249b | LOC_Os11g29840 | expressed protein |
| osa-miR6249b | LOC_Os11g31650 | expressed protein |
| osa-miR6249b | LOC_Os12g08360 | expressed protein |
| osa-miR6249b | LOC_Os12g29750 | protein of unknown function DUF502 domain containing protein, |
|  |  | expressed |
| osa-miR6249b | LOC_Os12g39310 | cytochrome P450, putative, expressed |
| osa-miR6249b | LOC_Os12g42280 | 9-cis-epoxycarotenoid dioxygenase 1, chloroplast precursor,<br>putative, expressed |

